# Aperiodic activity in the prefrontal cortex compensates for age-related decline in hippocampal theta and gamma oscillations during navigation

**DOI:** 10.1101/2024.10.03.616418

**Authors:** Sang-Eon Park, Thomas Donoghue, Seong Jin Lee, Chun Kee Chung, Joshua Jacobs, Sang Ah Lee

## Abstract

A critical gap remains in our understanding of age-related periodic and aperiodic electrophysiological changes across the human brain and their association with memory function. Using direct intracranial recordings from 69 epilepsy patients aged 19–61 performing a virtual navigation task, we separated oscillatory power from the aperiodic component. Hippocampal theta and gamma power decreased with age and correlated with impaired spatial memory. Age-related hippocampal spectral changes were replicated in an independent resting-state dataset. Aperiodic spectral slopes flattened in the prefrontal cortex but steepened in the hippocampus with age, suggesting region-specific changes in excitation–inhibition balance. Individuals with flatter dorsolateral prefrontal spectral slopes performed well despite reduced hippocampal theta and gamma power, suggesting a compensatory mechanism. This shift from hippocampal memory processes to prefrontal upregulation provides insight into cognitive resilience in aging and suggests potential targets for promoting healthy memory aging.

## Introduction

Despite the extensive evidence of navigational deficits in aging^1–3^ and Alzheimer’s disease^4–7^, the neurophysiological basis of individual differences in both age-related cognitive decline and maintenance in spatial memory has been difficult to pin down. To fully understand the many changes that occur in the brain across aging^8–14^, it is important to identify not only factors contributing to age-related memory decline but also the compensatory mechanisms that underlie individual cognitive performance^15^. Although past studies have identified alterations in neural activation related to both cognitive aging^3,5,16,17^ and compensation^11,18–21^, most thus far have been limited to indirect measurements of neural activity using functional neuroimaging^3,5,16–21^ or noninvasive electrophysiological recordings^22,23^. Some studies characterizing frequency-dependent oscillatory patterns based on source-localized MEG and EEG, have reported that spectral power decreases in low frequency oscillations (e.g. theta) but increases in the high frequency (i.e. beta and gamma) bands^22,23^. However, most of these studies measured neural activity during resting state and were therefore only indirectly associated with active memory processing. More recent task-based intracranial EEG (iEEG) studies provide greater insight into the functional neural correlates of human cognition^24^ and are critical to bridging insights on the neural basis of spatial memory between humans and nonhuman animals^25^.

The hippocampus, with its indispensable role in spatial cognition and memory, is particularly important due to its age-related degeneration^1–3^ and vulnerability to Alzheimer’s disease^4–6^. Given the primary role of hippocampal theta oscillations, often together with gamma oscillations^26^, in selective neural representations of spatial location^26^, distance^27,28^, and boundaries ^29,30^, its functional decline may induce age-related impairments in spatial cognition. Indeed, the recruitment of hippocampal theta in aged rats has been reported to be weaker during navigation compared to young rats^31,32^ and associated with a degeneration of hippocampal-dependent allocentric spatial memory^33^. Additionally, the reduction of theta oscillations in rodent models of AD^34,35^ has been associated with the accumulation of beta-amyloid^36^, cholinergic disruption^31,37^, and behaviorally observable memory deficits^38^. Surprisingly, however, despite the overwhelming evidence that hippocampal theta oscillation may play a crucial role in age-related cognitive decline in animal models, no studies thus far have shown direct intracranial recordings of such changes in human aging.

Interestingly, age-related decline in hippocampal integrity in humans^15,19^ is not enough to fully explain the high degree of individual variability in cognitive function across aging, particularly with respect to the maintenance of memory performance in the face of hippocampal decline. Some studies reported that extrahippocampal regions such as the prefrontal cortex are recruited for compensatory effects during memory encoding. For example, several neuroimaging studies reported that good-performing older adults exhibited increased dorsolateral prefrontal cortex (DLPFC) activation in the form of a posterior to anterior shift^39^ or hemispheric asymmetry reduction^40^. However, the exact nature of such prefrontal compensation and its relation to changes in hippocampal mechanisms are not known yet.

In characterizing the electrophysiological properties of the brain, it is becoming increasingly evident that separate analyses of the periodic and aperiodic components are important in order to avoid conflating different neurocognitive activities represented by each component^41–43^. In addition, dissociated periodic components have been reported to better represent their functional role in cognition and memory than conventional oscillatory power^44,45^. Furthermore, previous studies using ECoG, EEG, and MEG have reported a flatter slope of the power spectrum across the cortex in normal aging^46–49^ and its association with decreased cognitive function^48,50,51^. Recent studies suggest that this change in spectral slope may be related to an alteration in both local spiking frequency^41,52^ and excitation/inhibition balance^53^. Despite this, there is little work examining periodic and aperiodic activity in the human brain during the execution of a cognitive task, especially to identify their relationship to age-related memory decline^54^.

We set out to map both aperiodic and periodic electrophysiological markers of aging across the whole-brain (6935 electrodes) using iEEG recordings from 69 presurgical epilepsy patients (Fig. 1a) across a wide age range (19 to 61 years of age, median age 34), while they performed a computer-based spatial navigation task (Fig. 1b)^29,55,56^. In investigating individual variability and age-related degeneration in spatial cognition and memory, the hippocampus (221 electrodes) and DLPFC (117 electrodes) were our main regions of interest. We isolated periodic narrowband power from the aperiodic broadband component (spectral slope) using the specparam (spectral parameterization, formerly FOOOF) algorithm^57^ (Fig. 1c) to provide deeper insight into their age-related changes and association with cognitive decline.

**Figure 1.**
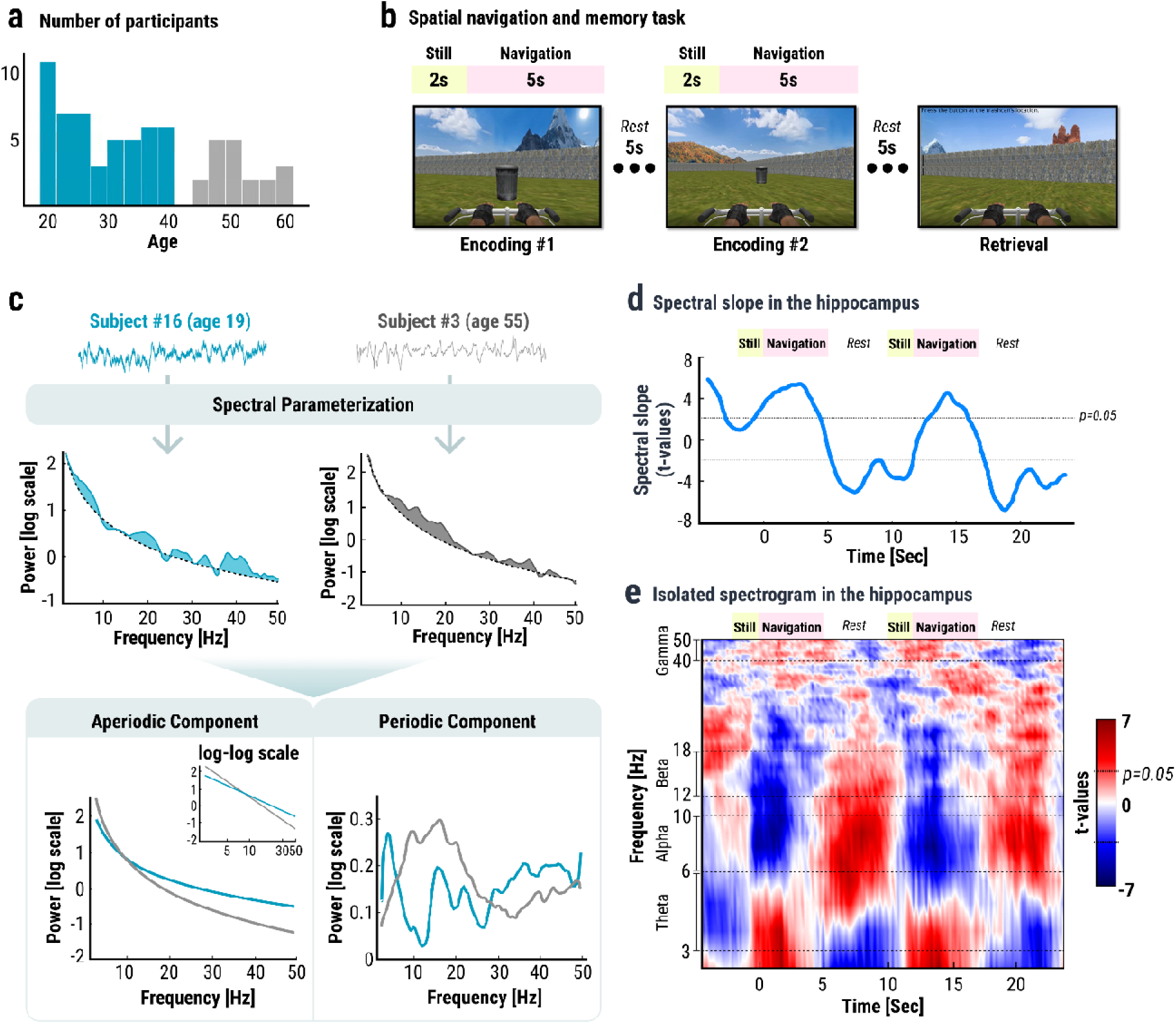
Overview of the spatial memory task and iEEG recordings. **a.** The distribution of age in years across our sample of 69 epilepsy patients. **b.** The spatial navigation task involving two passively-driven encoding phases, from different starting locations, to the same target location and a retrieval phase involving active-navigation from a new starting location. **c.** A schematic for calculating aperiodic and periodic neural components by applying spectral parameterization (*specparam*). An example of the power spectrum from a hippocampal electrode (middle row), showing the decomposed aperiodic (dotted line) and periodic components (shaded regions) during the encoding phase of the task. **d.** Time series of the hippocampal aperiodic component after smoothing during the encoding (still and navigation) and rest phases. **e.** Spectrogram of isolated periodic power in the hippocampus during the encoding (still and navigation) and rest phases.

## Results

### Age-related changes in aperiodic and periodic neural activity

To separately investigate the effects of aging on aperiodic and periodic (oscillatory) electrophysiological activity, we first isolated the two components from the iEEG signal during the spatial navigation task. Aperiodic spectral slope was measured by applying the *specparam* (spectral parameterization) algorithm^57^, and periodic narrowband power was calculated by subtracting the aperiodic spectral fit from the original log-power spectrum (Fig. 1c, see Methods for details). To test the engagement of each spectral component in spatial navigation, the aperiodic and periodic time series from hippocampal electrodes were calculated during the first and second encoding/rest periods and averaged across all trials (Fig. 1d and Supplementary Fig. 1c). We found a flatter aperiodic spectral slope during the encoding phase compared to the rest periods, which indicates that the hippocampus became more asynchronous and active (46–49) during navigation (Fig. 1d). Similar to previous studies^45,58,59^, theta and gamma power increased during the encoding phase compared to the rest periods, while alpha and beta power was suppressed (Fig. 1e).

We found two opposite age-related changes in spectral slope across brain regions (Fig. 2a, Supplementary Table 1, and see Supplementary Fig. 2 for the whole brain mapping): First, the spectral slope was flatter in older participants (i.e. positive correlation between a spectral slope and age) in the prefrontal, frontal, parietal, and occipital cortex, which is consistent with the interpretation that neural activity in the neocortical regions became more asynchronous and excitatory with age^46–49^. In contrast, a negative correlation between the spectral slope and age was observed in the temporal cortex and deeper brain regions. The steeper spectral slope in these regions may indicate a gradual transition into a synchronous and inhibitory neural state in aging. Notably, this suggests that the common finding of a flattening slope across age^46–49^ may be specific to the cortex (especially the PFC) and that the age-related changes in the hippocampal aperiodic activity may suggest its distinctiveness from the cortex.

**Figure 2.**
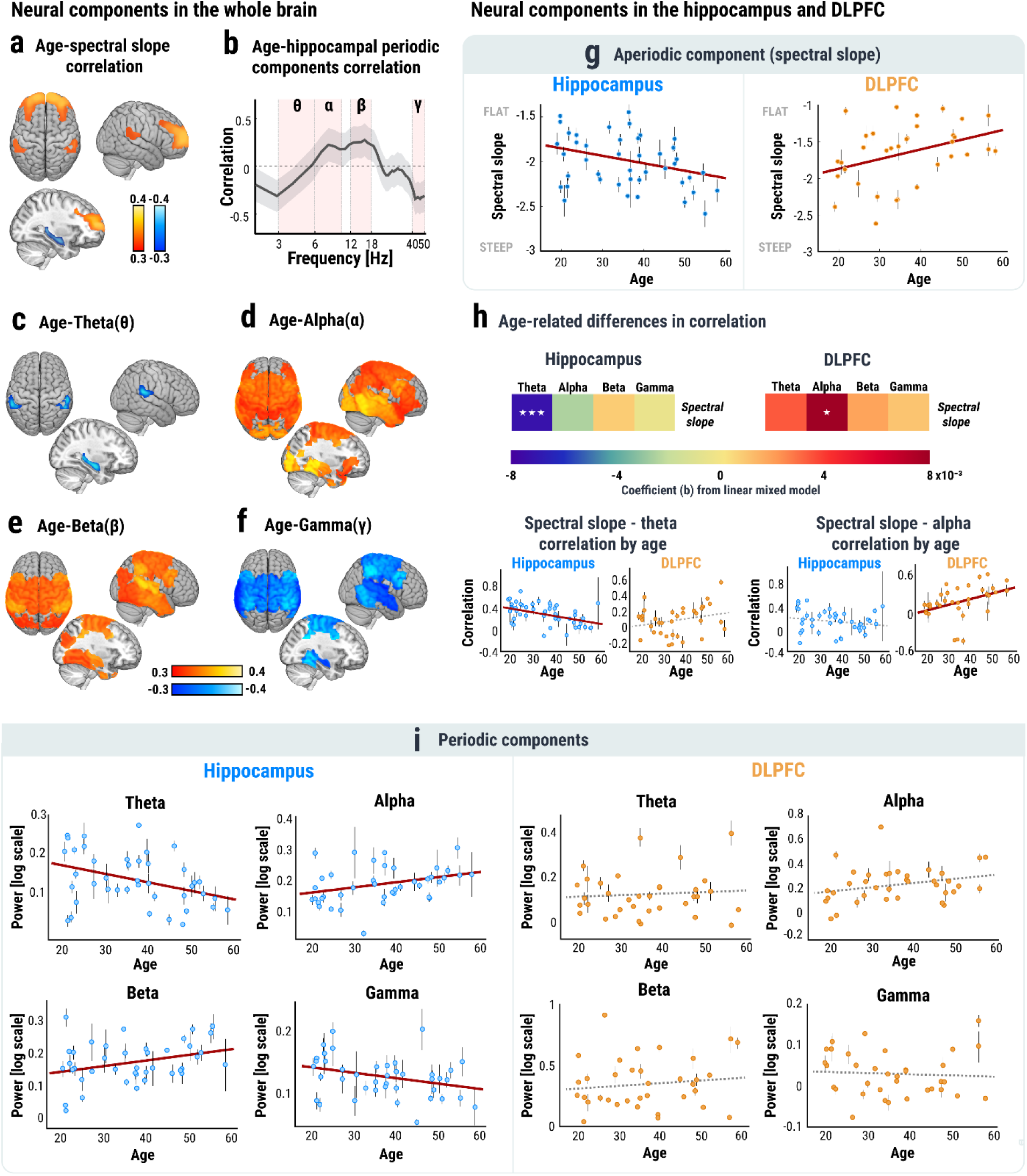
Age-related brain changes. **a.** A brain-wide map of Pearson correlations between age (ranges 19 to 61) and spectral slope during encoding. Only regions with a significant correlation (p<0.05) are indicated by the color bar. **b.** Correlation between age and power across frequency ranges between 2-50Hz in the hippocampus. **c-f.** A brain-wide map of significant correlations between age and theta (3-6Hz), alpha (6-10Hz), beta (12-18Hz), and gamma (40-50Hz) power. Only significant correlations (p<0.05) are highlighted and color-coded. **g.** A scatter plot of spectral slope across age in the hippocampus (left) and the DLPFC (right). **h.** Correlations between aperiodic and periodic components measured by trial-by-trial Pearson correlations. Color-coded values represent coefficients from a linear mixed-effects model with age as a predictor for the Pearson correlations between aperiodic and periodic components from all channels grouped by each participant. **i.** A scatter plot of periodic components during encoding across the whole age range for each frequency band. Each dot indicates the average value across multiple channels from one participant, and error bars indicate standard errors. Red lines indicate significance at p<0.05, while dotted black lines indicate that the predictor of the linear mixed-effects model was not significant. *** represents statistical significance at p<10^-3^, and * represents p<0.05.

We next compared the isolated periodic activity across aging. Correlations between age and power in each frequency range (theta: 3-6Hz, alpha: 6-10Hz, beta: 12-18Hz, and gamma: 40-50Hz; Fig. 2b, see Supplementary methods for details on the frequency range selection) showed that effects of aging on neural oscillations were frequency-dependent. (Fig. 2c-f and Supplementary Table 1). First, older adults showed significantly lower theta power in the hippocampus (Fig. 2c) (and superior temporal gyrus) but no other regions, revealing a unique age-related difference in hippocampal oscillations which was distinct from the cortex. Second, generally across the whole brain (with the exception of the PFC), both alpha and beta power was positively correlated with age (Fig. 2d and 2e), while gamma power was negatively correlated with age (Fig. 2f). These effects may together indicate a failure in coordination of inhibitory and excitatory neural functions^60–62^.

To verify that our results were not merely due to our specific sample of epilepsy patients, we performed a series of control analyses related to the individual seizure onset zone (SOZ). First, we found that age-related power differences in the hippocampus were preserved even when excluding electrodes from the SOZ (Supplementary Fig. 3). In addition, hippocampal theta from electrodes within the SOZ was not different from one outside of the SOZ, which indicated that in our sample hippocampal theta power was not significantly modified due to epilepsy.

Additionally, to ensure the robustness of the age-related pattern of electrophysiological activities in the face of other patient-specific differences, we analyzed an independent dataset comprising resting-state iEEG recordings from 46 bipolar electrode pairs implanted in the hippocampus of 27 epilepsy patients (ages 19-64, Supplementary Fig. 4-5). Linear mixed-effects models - with age, epilepsy duration, the number of anti-epileptic drug intake, or education length as a predictor, aperiodic/periodic components as dependent variables, and individual participants as a random-effect variable for grouping electrodes - confirmed our age-related patterns in hippocampal aperiodic/periodic activities (Supplementary Fig. 4 and Supplementary Table 3). We did not find any associations between hippocampal activity and disease duration, drug intake, or education (Supplementary Fig. 5 and Supplementary Table 4), with the single exception of a significant relationship between aperiodic spectral slope and epilepsy duration (b=-18.438×10^-3^, SE=6.814×10^-3^, p=0.01, Supplementary Fig. 5a). This relationship lost significance after controlling for age, but a trend still remained strong (r=-0.404, p=0.056, Supplementary Table 5) which suggested that a steeper spectral slope reflected the combined effects of aging and prolonged epilepsy.

### Contrasting patterns in age-related differences across the hippocampus and DLPFC

Of the above neural correlates, the most notable age-related changes occurred in the hippocampus and DLPFC, but in opposite directions from one another (Fig. 2g-i). By implementing linear mixed-effects models on neural components from these two brain regions, we first found that spectral slope during the encoding phase became steeper with age in the hippocampus (Fig. 2g: b=-7.672×10^-3^, SE=3.56×10^-3^, p=0.032) and flatter with age in the DLPFC (b=13.191×10^-3^, SE=5.539×10^-3^, p=0.019). No differences between the left and right hemispheres were found (Supplementary Fig. 6a and 6c).

In order to consider age-related changes in coordination among electrophysiological activities in the brain, we calculated correlations across the aperiodic and periodic components in a trial-by-trial manner and compared them across age (Fig. 2h). We found a significant interaction between age and region for predicting aperiodic-periodic correlation (linear mixed-effects model: age-region interaction in spectral slope-theta: F(1,334)=12.89, p<10^-3^; spectral slope-alpha: F(1,331)=14.691, p<10^-3^; spectral slope-beta: F(1,334)=4.017, p=0.046; spectral slope-gamma: F(1,332)=5.632, p=0.018), which indicated that the functional link between aperiodic and periodic components was weakened in the hippocampus but strengthened in the DLPFC. Specifically, older participants showed a significantly lower correlation between hippocampal theta power and spectral slope (Fig. 2h, age effect: b=-6.956×10^-3^, SE=1.966×10^-3^, p<10^-3^), suggesting a decrease in coordinated neural activities in the hippocampus. In comparison, the DLPFC in older participants showed a higher correlation between spectral slope and alpha power compared to the younger participants (Fig. 2h, age effect: b=8.228×10^-3^, SE=3.234×10^-3^, p=0.012). Therefore, spectral slope in the DLPFC not only became flatter with age (Fig. 2g) but also more correlated with other neural components.

#### Age-related differences in periodic oscillatory activity in the hippocampus

We found pronounced age-related oscillatory changes in the hippocampus (Fig. 2i): Along with a shifting of the peak frequency of the power spectrum from the theta band to higher alpha/beta band (Pearson correlation between age and the peak frequency: r=0.349, p=0.026), hippocampal theta and gamma power gradually decreased (Fig. 2i, age effect in theta: b= −2.123×10^-3^, SE=0.866×10^-3^, p=0.015; gamma: b=-1.029×10^-3^, SE=0.493×10^-3^, p=0.038) while alpha and beta power increased (alpha: b=2.632×10^-3^, SE=1.182×10^-3^, p=0.027; beta: b=1.902×10^-3^, SE=0.949×10^-3^, p=0.046) (see Supplementary Fig. 10 for analyses showing that these aging effects were weaker in other medial temporal lobe (MTL) regions). This trend across aging implies that the hippocampus became less active while simultaneously failing to maintain the neural activities required for successful memory performance (i.e. theta and gamma oscillations^44,45,58,59,61^). Among these differences, theta and gamma power decrease was specific to the left hippocampus (Interaction between age and region in theta: F(1,217) = 3.89, p=0.049; gamma: F(1,217)=4.723, p=0.031; See Supplementary Fig. 6 for details). Unlike the hippocampus, the DLPFC showed a similar oscillatory power pattern between the younger and older participants (Fig. 2i, age effect in theta: b=0.491×10^-3^, SE=1.505×10^-3^, p=0.745; alpha: b=3.468×10^-3^, SE=1.904×10^-3^, p=0.071; beta: b=2.54×10^-3^, SE=2.88×10^-3^, p=0.38; gamma: b= −0.3×10^-3^, SE=0.751×10^-3^, p=0.69). Interestingly, these neural markers of aging in the hippocampus and DLPFC were not found using the original power spectrum (i.e. before separating the oscillatory activity from aperiodic component, Supplementary Fig. 7). In addition, prior EEG and MEG studies reported age-related decreases in alpha and increases in gamma power^22,23,63^, whereas we observed the opposite. This discrepancy may result from the inclusion of the broadband aperiodic spectral slope, which can interfere with conventionally measured band power^44^, underscoring the need to separate these components to accurately capture electrophysiological changes in aging.

The above findings together raise the possibility that age-related functional degeneration of the hippocampus during navigational encoding may emerge in parallel with a functional reorganization of the DLPFC. If this hypothesis is true, then these age-related differences in brain activity should be reflected in the spatial memory performance of older participants.

### Age-related spatial memory deficits and their hippocampal neural correlates

Following the two encoding phases, in which participants were passively driven to a target object from two different starting positions in an open maze, they freely navigated from a random location to the hidden object and responded by pressing a button (retrieval, Fig. 1b). To assess spatial memory performance, distance errors were calculated by computing the distance between the reported response and the actual target location (Supplementary Fig. 8a). General linear models were implemented to characterize age-related differences in behavioral measures with the observation weights (negatively) proportional to the age-dependent density of samples to balance out the smaller sample size of older participants (Supplementary Fig. 8b)^64,65^. The older participants performed worse compared to the younger participants, with higher distance error (Supplementary Fig. 8c, age effect: b=0.111, SE=0.051, p=0.034) and longer travel time to their destination (Supplementary Fig. 8d, b=0.296, SE=0.097, p=0.003).

To find the neural correlates linking aging and spatial memory degeneration, we implemented linear mixed-effects models with distance error as a predictor, aperiodic/periodic components during encoding from each electrode in the hippocampus and DLPFC as dependent variables, and individual participants as a random-effect variable for grouping electrodes. Hippocampal theta and gamma power were tightly associated with distance error (Fig. 3a and 3b theta: b=-3.853×10^-3^, SE=1.89 ×10^-3^, p=0.043; gamma: b=-2.702×10^-3^, SE=1.047×10^-3^, p=0.011), while other periodic band power (i.e. alpha and beta) and the aperiodic spectral slope did not show a difference across distance error (Supplementary Fig. 9a, spectral slope: b=-8.695×10^-3^, SE=7.856×10^-3^, p=0.27; alpha: b=-2.304×10^-3^, SE=2.656×10^-3^, p=0.387; beta: b=3.92×10^-3^, SE=2.026×10^-3^, p=0.054).

**Figure 3.**
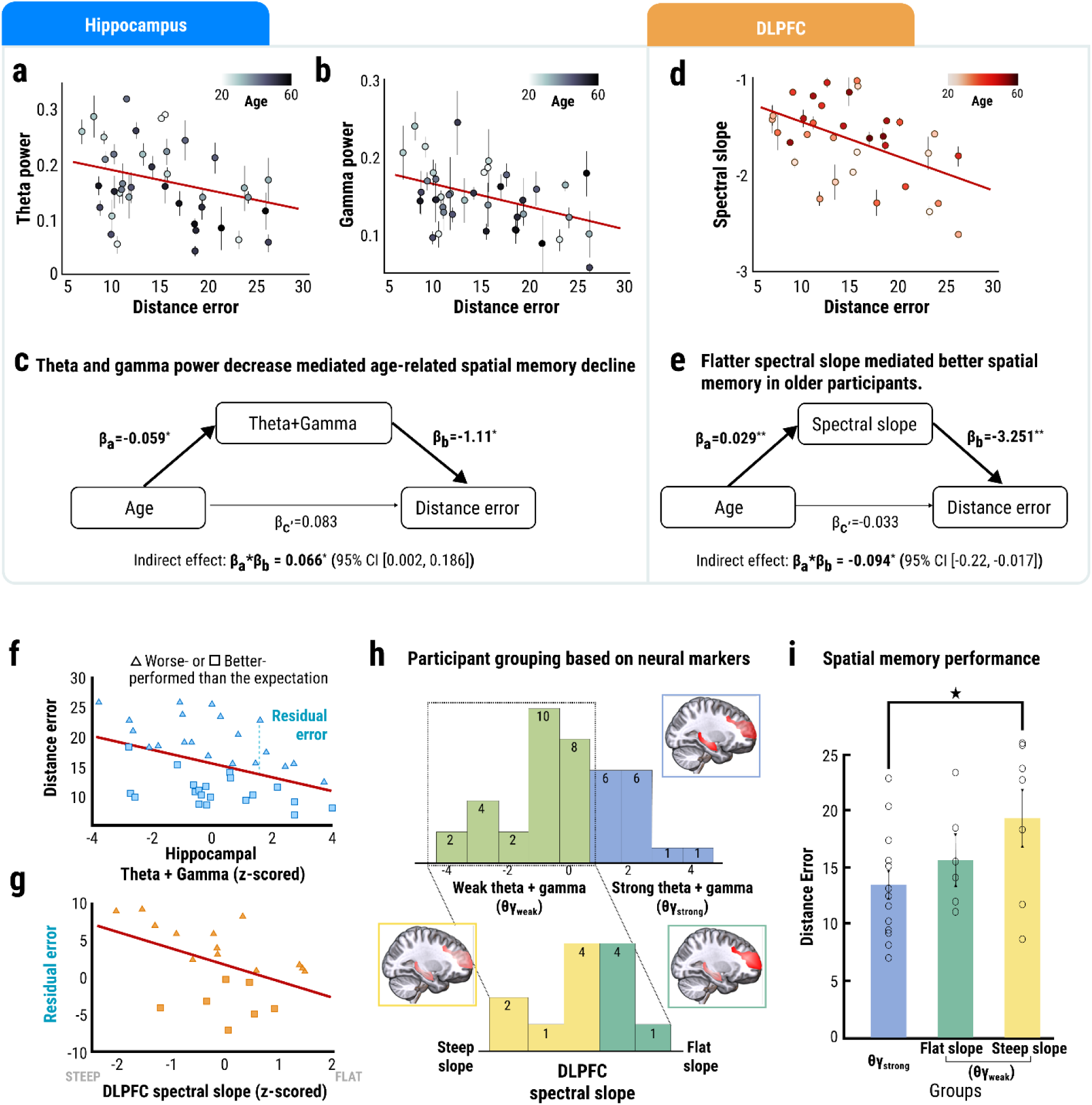
Neural correlates of age-related spatial memory performance differences. **a-b.** Scatter plots of (**a**) hippocampal theta and (**b**) gamma power across the whole distance error range. Each dot indicates the average value across multiple channels from one participant, and error bars indicate standard error. Color-coded values represent the age of the corresponding participants. **c.** A mediation analysis showed that an age-related decrease in the sum of z-scored theta and gamma power mediated distance error increase. **d.** A scatter plot of spectral slope in DLPFC and distance error. Each dot indicates the average value across multiple channels from one participant, and error bars indicate standard error. **e.** A result of mediation analysis showed that a flatter spectral slope in the DLPFC mediated reduced distance error in older participants. **f-i. Hippocampal dysfunction compensated by DLPFC spectral slope. f.** A scatter plot showing a linear relationship between hippocampal theta + gamma power and distance error. The *residual error* was defined as the deviance in distance error from the expected value based on the sum of z-scored hippocampal theta and gamma powers. Triangles indicate participants who performed worse than the expectation; rectangles indicate participants who performed better than the expectation. **g.** A scatter plot showing a linear relationship between the DLPFC spectral slope and the *residual error*. The flatness of the DLPFC spectral slope was correlated with spatial memory performance. **h.** The number of participants after grouping based on hippocampal theta + gamma power (top, θγ_strong_ and θγ_weak_); subsequent grouping of the θγ_weak_ participants based on DLPFC spectral slope (bottom). The number inside the bars indicates the number of participants **i.** Distance error from the spatial memory task across the three groups, showing worst performance in participants with weak theta + gamma and steep spectral slope in DLPFC. **a, b, d, f, g.** The red lines indicate a significant level at p<0.05. ** indicates a significant level at p<0.01. * indicates p<0.05.

The relationship between theta and gamma power and performance was pronounced in the hippocampus compared to other MTL regions (i.e. entorhinal cortex, perirhinal cortex, and amygdala; region-distance error interaction in theta: F(3,440)=3.657, p=0.013; gamma: F(3,444)=3.681, p=0.012; Supplementary Fig. 10f and 10g) and stronger in the left hippocampus compared to the right (region-distance error interaction in theta: F(1,217) = 7.479, p=0.007; gamma: F(1,217)=5.269, p=0.023; Supplementary Fig. 6b). Lastly, a mediation analysis suggested that decreased hippocampal theta and gamma power significantly contributed to age-related spatial memory decline (Fig. 3c, indirect effect = 0.066, 95% CI [0.002, 0.186], p=0.046).

Complementary to the findings on hippocampal theta and gamma power, the spectral slope in the DLPFC was flatter in participants with lower distance error (Fig. 3d, b=-0.035, SE=0.011, p=0.003). Additionally, a mediation analysis indicated that the age-related flattening of the spectral slope in the DLPFC (Fig. 2g) contributed to better performance (Fig. 3e, indirect effect: −0.094, 95% CI [−0.22, −0.017], p=0.024), raising the possibility that a compensatory over-recruitment of the DLPFC aids in the maintenance of spatial memory ability in aging. In contrast, the periodic neural features (i.e. theta, alpha, beta, and gamma) in the DLPFC were not associated with spatial memory performance (Supplementary Fig. 9b).

### Spectral slope in the DLPFC compensates for hippocampal dysfunction

Further evidence of compensation was revealed by a tight association between the DLPFC spectral slope with residuals from linear regression model of distance error using the hippocampal markers (Fig. 3g, Pearson’s correlation r=-0.455, p=0.044). Upon closer look at the negative linear relationship between distance error and the sum of z-scored hippocampal theta and gamma power (Fig. 3f, Pearson correlation r=-0.394, p=0.012), the subset of individuals who performed better than expected given their hippocampal theta and gamma power (i.e. negative residuals, Fig. 3g and 3f) showed increased activation in the DLPFC as inferred from a flatter spectral slope. On the contrary, the subset of participants who performed worse than expected from their hippocampal theta and gamma power (i.e. positive residuals, Fig. 3g and 3f) did not exhibit an upregulation of the DLPFC, as inferred by a steeper DLPFC spectral slope.

To test the possibility that increased DLPFC excitability may compensate for weak hippocampal function (11, 18–21), we first divided the participants according to their hippocampal theta and gamma power. The strong theta + gamma group (θγ_strong_) (Fig. 3h) consisted of the top one-third of the participants (n=14; age: mean=28.962, SE=2.411), and the weak theta + gamma group (θγ_weak_) consisted of the bottom two-thirds (26 participants). The θγ_weak_ group was further divided into two groups based on their DLPFC spectral slope (“steep slope” and “flat slope” groups, see Fig. 3h bottom). Average distance error was lowest in the θγ_strong_ group, as expected (Fig. 3i). Of the two θγ_weak_ groups, the steep slope group (7 participants; age: mean=38.519, SE=3.446) performed significantly worse than the θγ_strong_ group (t(19)=2.334, p=0.03) but the flat slope group (6 participants; age: mean=36.491, SE=4.248) showed comparable performance with the θγ_strong_ group (t(18)=1.02, p=0.321). These results are consistent with the interpretation that the over-recruitment of the DLPFC serves to compensate for weak hippocampal oscillatory power and may reflect individual cognitive resilience to hippocampal aging.

### Compromised hippocampal theta correlate of scene-based navigation underlies impaired memory retrieval

We next asked whether the changes in hippocampal activity across age can be associated with changes in spatial cognitive processes involved in navigation. Given previous results showing that high-performing participants in this task oriented towards a similar view (i.e., scene) from the encoding phase^55^, we categorized the trials into *Match* and *Non-match* conditions, based on the overlap in scenes between the encoding and retrieval paths (Fig. 4a and 4b, see Methods and Supplementary Fig. 11 for details). As expected, accuracy was higher for the *Match* condition compared to the *Non-match* condition (t(65)=9.593, p<10^-3^); this difference was higher in younger than older participants (Fig. 4c, RMANOVA interaction between age and condition: F(1,64)=11.439, p=0.001), with a significant difference across aging only in the *Match* condition (Supplementary Fig. 11c, Pearson correlation between age and distance error for *Match*: r=0.374, p=0.002; Nonmatch r=-0.146, p=0.239). A linear mixed-effects model with age and scene condition (i.e. *Match* or *Nonmatch*) as a predictor, neural components as dependent variables, and individual participant as a random-effects variable showed a main effects of condition (F(1,429)=11.991, p<10^-3^), a main effect of age (b=-1.831×10^-3^, SE=0.872×10^-3^, p=0.036), and an interaction between the two, indicating that age-related theta power decline was specific to the *Match* condition (Fig. 4d, F(1, 429)=7.423, p=0.007; *Match*: beta=-1.797×10^-3^, SE=0.881×10^-3^, p=0.043; *Nonmatch*: beta=-0.275×10^-3^, SE=0.841×10^-3^, p=0.744). These results suggested that reduced theta engagement in the Match condition may contribute to impaired scene-based memory retrieval in older participants. Match–Nonmatch differences in spectral slope, alpha, beta, and gamma power were not associated with age (Supplementary Fig. 11h).

**Figure 4.**
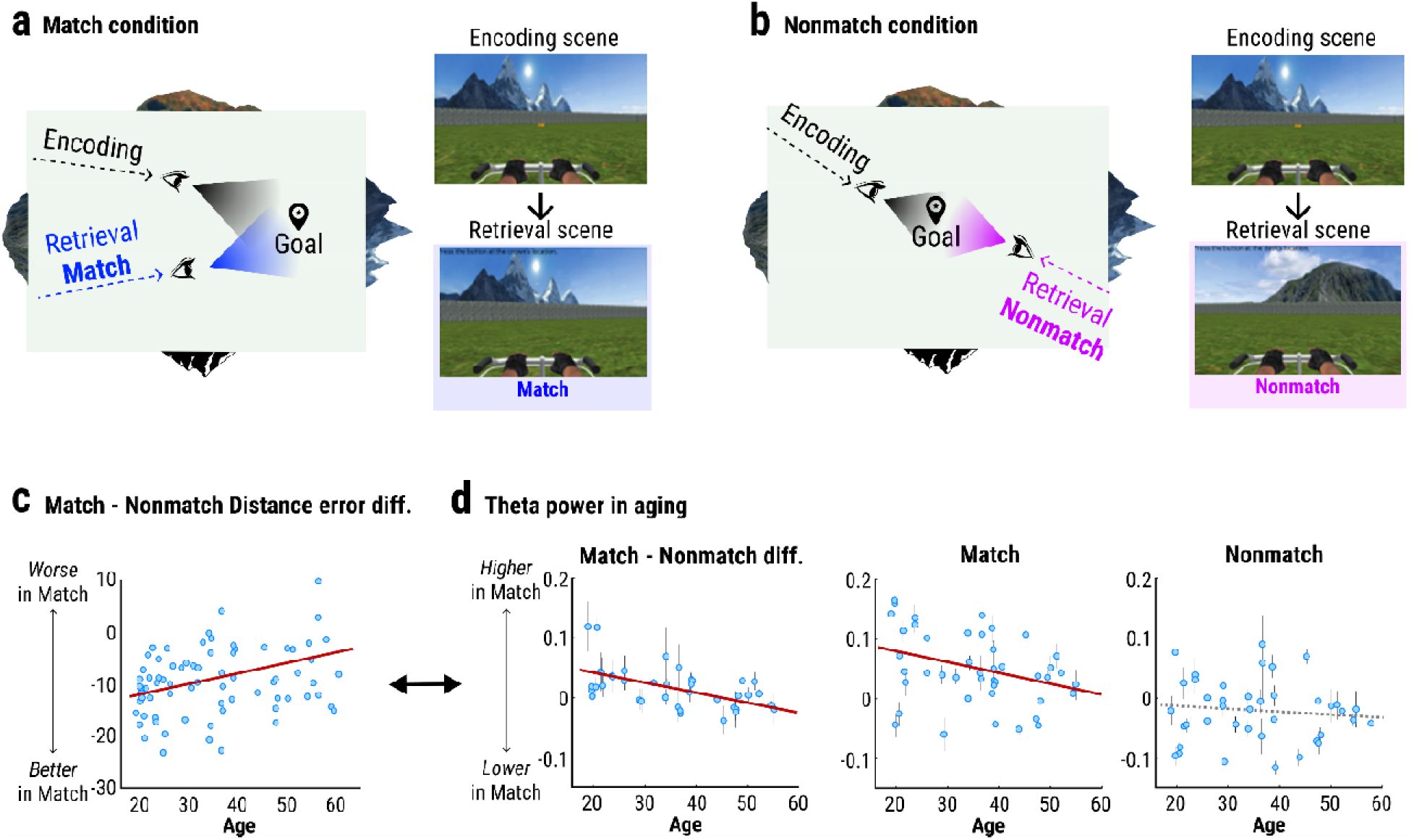
Impaired scene utilization during retrieval in aging and its neural correlates. **a-b.** Left: A bird-eye view schematic of the virtual environment with an example of encoding and retrieval paths for *Match* (**a**) and *Nonmatch* (**b**) trials. Right: Examples of landmark views during encoding and retrieval across the two conditions. **c.** *Match* - *Nonmatch* error difference across the whole age range. **d.** Hippocampal theta power in the *Match* (middle) and *Nonmatch* (right) conditions and their difference (*Match* - *Nonmatch*; left) across age. Each dot indicates the average value across multiple channels from one participant, and error bars indicate standard error. The red lines indicate a significance level at p<0.05 of the age effect, based on the linear mixed-effects models.

## Discussion

In this study, we investigated the aperiodic and periodic intracranial electrophysiological markers of aging throughout the brain, as well as the relationships among them, and identified those that may be critical in determining individual age-related decline and compensation in spatial memory. As expected from the previously established role of hippocampal oscillations in spatial memory, a decrease in theta and gamma power during the spatial navigation task was significantly associated with age-related memory decline. In contrast, we found a potential compensatory mechanism in the DLPFC as indicated by the flatter spectral slope in individuals with good performance despite their lower hippocampal function.

Our results showed that older participants were impaired in their ability to navigate based on the matched spatial view of the target location (between encoding and retrieval). Therefore, spatial memory performance in our task was dependent on the ability of the participants to use spatial cues such as the boundaries and distal landmarks. Given the role of theta oscillations in processing of landmarks^66–68^ and spatial boundaries^29,30,69^, the older adults’ failure to engage hippocampal theta in this task may underlie their difficulty utilizing spatial cues to form a cognitive map.

What could be the underlying neurophysiological mechanism of the decrease in theta and gamma power? Previous studies suggested that cholinergic projections from the basal forebrain to the hippocampus degrades with aging^70^. Interestingly, pharmacological enhancement of cholinergic responses, such as the administration of cholinergic agonist carbachol, reversed hippocampal place-coding deficits and theta power decreases in aged rats during novel maze exploration^31,37^. In contrast, administration of cholinergic-antagonist scopolamine induced memory impairments in younger adult humans that were similar to those observed in aged individuals^71^. Similarly, in a recent study, reduced theta power was observed following the administration of scopolamine in epilepsy patients^72^. Additionally, other rodent studies showed that the cholinergic blockade reduced coordinated activities between theta and gamma oscillations^73^. The evidence as a whole suggests that hippocampal theta and gamma power decrease and spatial memory degeneration are highly associated with changes in cholinergic processes in the brain.

Our interpretation of the flatter spectral slope of the DLPFC in high-performing older participants as a compensatory marker draws upon previous fMRI studies reporting increased activation of the DLPFC in older adults to compensate for age-related memory decline^39,74^. In addition, the age-related association between spectral slope and alpha power raises the possibility of a functional neural reorganization to enhance prefrontal cognitive processes for maintaining spatial memory performance in spite of a degenerated hippocampus.

While there are numerous reports of a flattening of the spectral slope across age in the cortex^46–49^, there has been surprisingly little investigation of aperiodic electrophysiological changes in deeper brain regions ^47^. For the first time using direct intracranial recordings, we have revealed the opposite pattern of a steepening of the spectral slope across aging in the hippocampus. These findings converge upon a recent analysis of source-projected MEG data, which reported that older subjects had steeper slopes in deep structures including the hippocampus^75^. Overall, these results emphasize the importance of a whole-brain approach to characterizing aperiodic activity, particularly in the hippocampus, for understanding specific electrophysiological changes in the aging brain.

The steeper spectral slope and theta/gamma power decrease in the hippocampus contrasted by a flatter spectral slope without oscillatory changes in the DLPFC may be attributed to differential changes in these areas at the cellular level. Transmission of the neocortical inhibitory neurotransmitter GABA has been reported to degrade in aging^76,77^, which may result in an increased overall activity level, as indicated by a flatter spectral slope in older individuals. Decreased GABA signaling may also have contributed to the gamma power decrease in our study, given previous findings in AD models of rodents^78^. Contrary to reports of disrupted GABA signaling in the hippocampus^79^, we observed a steeper spectral slope typically associated with an increased inhibition, potentially caused by anatomical or functional changes in synapses and reduction in excitatory postsynaptic potentials (EPSP)^80^. Decreases in NMDA receptors in aging may cause a similar reduction in long-term potentiation (LTP)^81^. Other possible mechanisms for lower levels of neural activities include increased calcium channel conductance and higher after-hyperpolarizing potentials^81^ and decreased excitatory inputs to the hippocampus (e.g., due to changes in dopamine, norepinephrine, acetylcholine)^82^.

By leveraging the advantages of task-based iEEG data from a large sample of participants, we not only provided an overview of the electrophysiological changes in the aging human brain but also discovered a shift in the electrophysiological activity during navigation from the hippocampal theta oscillations to aperiodic DLPFC excitation in compensating for the age-related decline in memory function. Our findings suggest that older adults who maintain a high level of performance may switch from their reliance on precise hippocampal spatial mapping to a more efficient information processing strategy via domain general prefrontal regulatory mechanisms^83,84^. Future studies identifying the neurobiological origins of these cognitive changes may open new doors to the development of targeted neuromodulatory interventions and prevention methods for maximizing cognitive reserve in aging.

## Methods

### Participants

For the main analyses, 69 presurgical epilepsy patients (30 male and 39 female) between ages 19 and 61 (Fig. 1a) performed a virtual navigation task on a laptop computer while intracranial EEG (iEEG) signals from a total of 6935 electrodes were recorded. This study was approved by local institutional review board protocols at eight testing sites (Thomas Jefferson University Hospital (Philadelphia, PA); University of Texas Southwestern Medical Center (Dallas, TX); Emory University Hospital (Atlanta, GA); Dartmouth-Hitchcock Medical Center (Lebanon, NH); Hospital of the University of Pennsylvania (Philadelphia, PA); Mayo Clinic (Rochester, MN); National Institutes of Health (Bethesda, MD); and Columbia University Hospital (New York, NY)), as well as the institutional review board of the University of Pennsylvania (data coordination site) and the Human Research Protection Office at the Space and Naval Warfare Systems Command Systems Center Pacific. All participating patients provided informed consent. The raw data are available online (http://memory.psych.upenn.edu/) and were used in previous studies ^29,55,56^, but the results presented here are novel.

For the control analyses, 27 epilepsy patients between ages 19 and 64 were recruited and resting-state iEEG signals were recorded from 46 electrodes in the hippocampus. This study was approved by local institutional review board protocols at Seoul National University Hospital (Seoul, Republic of Korea). All participating patients provided informed consent.

### Spatial memory task

Each encoding phase (2 encoding phases per trial x 48 trials per session, 1–6 sessions per participant) included a 2s period during which participants were presented with a still scene from the starting point in the environment followed by a 5s period of passive navigation toward a target object. At the start of the encoding phase, the target object appeared in the environment, and participants were rotated (1s) and driven (3s) toward the target with a constant speed and then stopped (1s) at the target location. A black screen appeared for 5s, and then the encoding phase was repeated a second time from a different starting location (to the same target location as the first encoding phase). A black screen appeared again for 5s, and then the retrieval period began, in which participants were placed in a random starting location in the environment from which they had to freely navigate to the hidden target object using a joystick and press a button to mark their response. Participants then received feedback on their performance via an overhead schematic of the environment showing the actual location of the target object and their response. Distance error was the main metric for performance evaluation and represented how far the participants’ response was from the actual target location.

### Intracranial recordings and electrode localization

iEEG was recorded at a sampling rate of 500 Hz or above. Bipolar referencing was applied by computing the voltage difference between pairs of adjacent electrodes to reduce non-physiological artifacts. The anatomical location of each bipolar pair was labeled as the midpoint between the two electrodes. Lastly, the raw iEEG data were notch-filtered using a Butterworth filter at 60 Hz.

We determined the location of each electrode by co-registering a post-surgical CT scan to T1-weighted structural MRIs (0.5 × 0.5 × 2 mm resolution) taken prior to implantation. First, electrode localization in cortical regions was determined by Freesurfer’s automated cortical parcellation based on the Desikan-Killiany brain atlas^85^. Second, depth electrodes in the MTL were labeled by using a multi-atlas-based segmentation technique^86,87^ followed by a visual inspection from a neuroradiologist on post-implant CT scans, co-registered with the MRI scans.

### Calculating aperiodic and periodic neural components

We applied the *specparam* tool (v1.0, formerly *FOOOF*) to parameterize periodic and aperiodic components from neural power spectra (see Donoghue et al., 2020 for methodological details). *Specparam* quantifies the aperiodic (1/f-like) component, equivalent to the slope of log frequency-log power spectrum, and then detects and quantifies putative oscillations by calculating the area of power above this aperiodic component. Power spectra during the encoding phase were calculated by using Welch’s method (1.6s moving windows, 0.8s overlap) separately for each trial. Spectral parameterization was applied to the frequency range of 2-50 Hz, in ‘fixed’ aperiodic mode (fitting a single slope value, without a ‘knee’ or bend in the aperiodic component) and with other settings (peak width limits between 0.5 to 12 Hz, minimum peak height of 0, peak threshold of 2, and unlimited maximum number of peaks). The frequency range was chosen to fit a range that can be modelled without a knee that is often present in intracranial recordings. The absence of a knee was visually confirmed from average power spectra. This method also provides measures of model fit quality, by measuring the residual errors and R-square values between the full model fit and the original power spectrum. By implementing a linear mixed-effects model with age as a predictor, we confirmed that the residual errors were not different across ages in both regions (hippocampus: b=0.09×10^-3^, SE=0.353×10^-3^, p=0.8; DLPFC: b=0.665×10^-3^, SE=0.476×10^-3^, p=0.165). R-square values were high for all participants and regions (mean±SE in the hippocampus: 0.984±0.006; DLPFC: 0.979±0.011).

To test neural engagement in the spatial memory task, we measured the temporal pattern of both aperiodic and periodic components across two encoding and rest phases (Fig. 1d and 1e). Power spectra were calculated with a 3s moving window (step size=0.2s), and spectral parameterization was applied to the frequency range of 2-50 Hz with a frequency resolution of 0.5 Hz. Aperiodic and periodic components at every time point in each participant were z-scored for every participant having the same mean and standard deviation of the time series. Then, t-tests were applied for neural components at every time point at a participant level to compare with the zero mean.

### A characterization of aging effects across the whole brain

We investigated age-related differences across neocortical regions divided as Brodmann areas (BA1&2&3&5 (261), BA4&6 (663), BA7 (173), BA8 (168), BA9 (291), BA10 (263), BA11 (272), BA13 (299), BA18 (122), BA19 (293), BA20 (433), BA21 (832), BA22 (373), BA31 (172), BA32 (141), BA37 (204), BA38 (204), BA39 (192), BA40 (417), BA41&42 (119), BA44 (65), BA45 (112), BA46 (117), BA47 (192); the numbers inside the parentheses indicate the number of bipolar-paired electrodes) and deep brain regions such as the hippocampal formation and its subregions (i.e. the CA1 (121), CA3 & Dentate gyrus (65), and subiculum (28)), as well as the amygdala (142), entorhinal cortex (26), perirhinal cortex (63), and striatum (148). The number of participants in each region is indicated in Supplementary Table 2. Pearson correlations between age and neural components were measured at the participant level after averaging across all channels within the region. Outliers more than three scaled median absolute deviation (MAD) away from the median value were removed at the trial, channel, and participant levels.

### Aging effects in the hippocampus and DLPFC

We characterized age-related differences in neural components, both at the participant and trial levels. For the participant-level analysis, we implemented linear mixed-effects models with age as a predictor, neural components from all channels as a dependent variable, and individual participant as a random-effects variable for grouping channels^88,89^. For the trial-level analysis, we quantified the coupling between two neural components by calculating Pearson correlation coefficients between neural components across trials and then compared across age by implementing linear mixed-effects models. Outliers more than three scaled MAD away from the median value were removed at the trial and participant levels.

### Neural markers associated with spatial memory performance

We identified age-related neural components associated with spatial memory decline or compensation in the hippocampus and DLPFC. Specifically, linear mixed-effects models were implemented with distance error as a fixed-effect predictor, neural components from all channels as a dependent variable, and individual participant as a random-effect grouping variable.

Next, mediation analyses were performed to find neural markers of age-related spatial memory decline: distance error from each participant was the dependent variable, the mediator was a neural component (either hippocampal theta + gamma power or DLPFC spectral slope), and age was the independent variable. Each neural component was z-scored at the participant level. We tested whether there was a significant indirect (mediation) effect (product of coefficients = a*b from the equation (1) below) for each neural component using the bootstrapping method by random resampling of original data with replacement^90^ (repeated 10000 times), along with the remaining direct effect between age and performance by c’ from the equation (2).

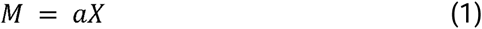

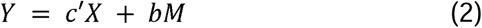

### DLPFC spectral slope compensates hippocampal theta and gamma dysfunction

A general linear model was implemented on distance error with the sum of hippocampal theta and gamma power (each z-scored at the participant level). The residual distance error from the model was then used as a dependent variable for another general linear model with DLPFC spectral slope (z-scored at the participant level) as a predictor. In addition, we divided participants into three groups based on their neural activities. The first grouping was based on hippocampal theta and gamma power: The strong theta + gamma (θγ_strong_) group consisted of participants who showed top third (N=14) of a sum of hippocampal theta and gamma power, and the weak theta + gamma (θγ_weak_) group were the other remaining participants. The θγ_weak_ group was divided into two groups based on whether their spectral slope was above or below the average (“Steep slope” and “Flat slope” groups). Only participants with electrodes in both the hippocampus and DLPFC were included in the subsequent subdivision of the weak group. Lastly, distance error in the spatial memory task was compared across the three groups (θγ_strong_ group, Steep slope group, and Flat slope group) by applying two-sample t-tests.

### Comparison of scene-dependency across age during retrieval

We divided each trial into Match/Nonmatch conditions based on the paths traveled during the two encoding and retrieval phases to assess the participants’ utilization of scenes for spatial mapping. First, to isolate the cases in which the participants were provided with a particular scene associated with the target location, we selected the trials in which the angle difference between the two encodings paths was less than 90 degrees. From there, the Match condition consisted of those trials in which the participants’ retrieval direction was less than 30 degrees offset from to one of the two encoding paths, and the Nonmatch condition consisted of those trials that in which the participants’ retrieval direction was higher than 55 degrees from either of the two encoding paths. The choice of the two criteria (i.e. 30 and 55 degrees) were verified through behavioral analysis that showed that the performance was clearly different between two conditions and that a ‘transition zone’ existed between the two angles (Supplementary Fig. 11d).

### Comparison of neural components between the Match and Nonmatch conditions

Hippocampal correlates of retrieval in the Match condition were searched and compared across the entire age range. Free navigation yielded heterogeneous behavioral patterns across subjects and trials (e.g. varying duration for still and navigation periods), so we focused on the navigation onset (3s) which we hypothesized to be the most critical period for utilizing the scene and deciding the direction of one’s trajectory to the target location. We implemented linear mixed-effects models with age and condition as a predictor, neural components as a dependent variable, and participants as a random-effect grouping variable.

## ACKNOWLEDGMENTS

This study was supported by Creative-Pioneering Researchers Program through Seoul National University and the National Research Foundation of Korea (RS-2024-00342897, 2021M3E5D2A01023891, and 2020K1A3A1A19088932).

## Data Availability

The raw data are available online (http://memory.psych.upenn.edu/).

## Code Availability

Requests for the codes can be sent to the corresponding author.

## Author Contributions

S-E.P., J.J., S.A.L., conceived and designed the study. J.J. and S.A.L. collected the behavioral and iEEG data. S.J.L. and C.K.C collected Korean iEEG data. S-E.P, T.D., J.J., and S.A.L. analyzed the data, prepared the figures, and wrote the main manuscript text. J.J. and S.A.L. supervised the study and revised the manuscript. All authors reviewed and approved the final manuscript.

