## Supplementary Information for "Aperiodic activity in the prefrontal cortex compensates for age-related decline in hippocampal theta and gamma oscillations during navigation"

**Supplementary Methods**

The frequency band was defined based on a data-driven approach. Isolated band power was calculated and averaged into 1Hz bins. We calculated the correlation between age and band power from the hippocampus at each frequency bin and found that the sign of the correlation changed around 6 and 18Hz. Additionally, we observed that the correlation at 10-12 Hz was marginal compared to the surrounding frequency bands. Based on these observations, we defined our theta (3-6Hz), alpha (6-10Hz), and beta (12-18Hz) bands. The dissociable pattern between 3-6 and 6-10 and 12-18 Hz was similar to what was observed in Rudoler et al 2023. The correlation was marginal between 18-40 Hz and became significant again after 40Hz, so we defined the gamma band as 40-50Hz.

The pattern from the isolated periodic band power was different from the one observed with the raw power spectrum. Specifically, some studies reported that the raw spectral power of the canonical alpha band (8-12Hz) decreased with age, which is opposite from what was observed in this study. We suspect that the alpha power decrease was a misinterpretation of flatter spectral slope in the aging cortex.

**Supplementary Figure. 1 Temporal dynamics of neural features during spatial encoding**

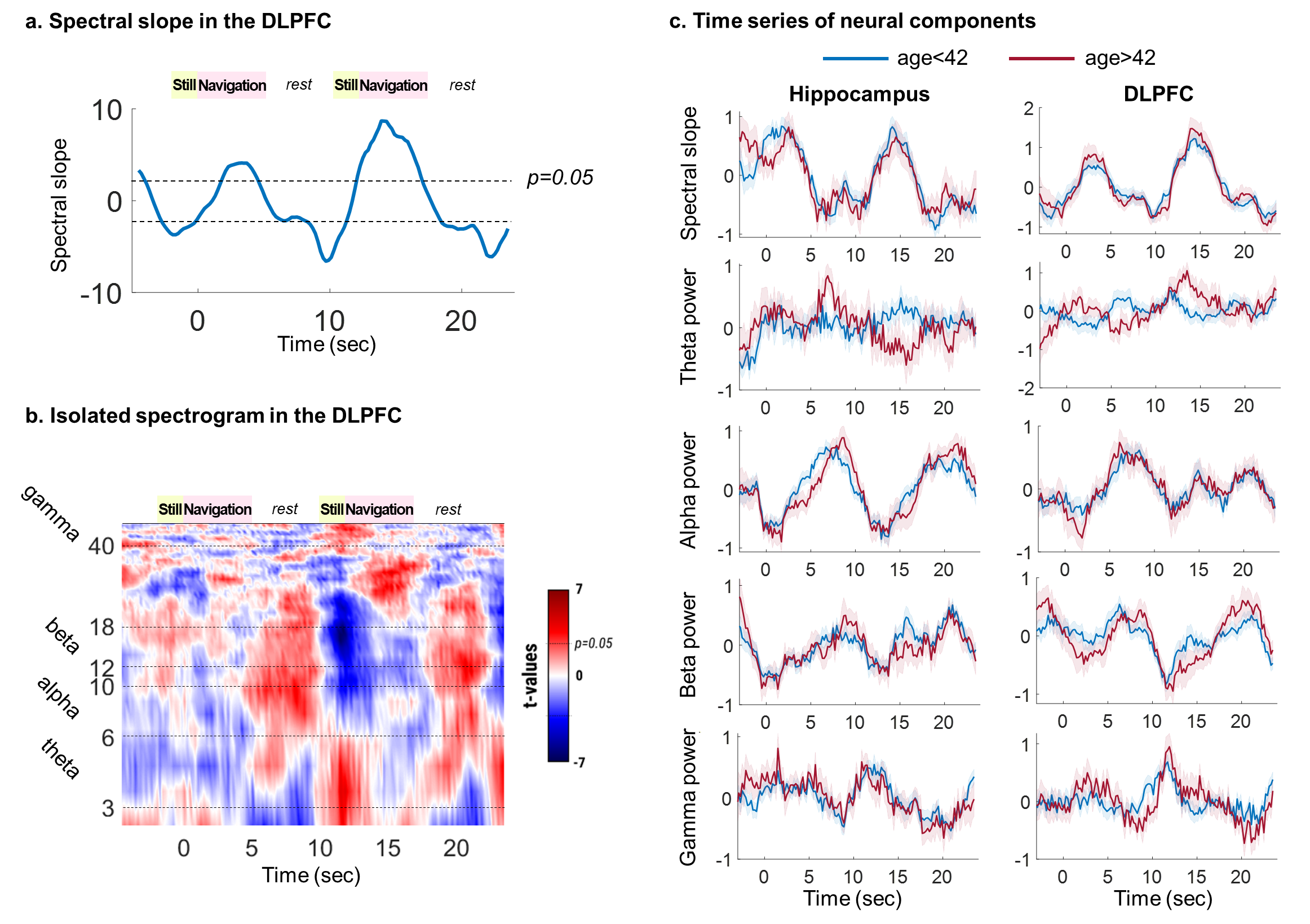

**a-b.** We calculated temporal dynamics (window size=3s and moving step=0.2s) of aperiodic and periodic features in the DLPFC across two encoding and rest phases. Aperiodic and periodic components at every time point in each participant were z-scored for every participant having the same mean and standard deviation of the time series. Then, t-tests were applied for neural components at every time point at a participant level to compare with the zero mean. Theta and gamma power in the DLPFC showed different patterns from the hippocampus (Fig. 1d and 1e) especially during the first encoding phase, while spectral slope, alpha, and beta power showed a similar pattern. **c.** Time series of neural components during the encoding (i.e. still and navigation) and rest phases and its comparison between younger and older participants (grouped by age over or under 42 for visualization purposes).

**Supplementary Figure 2. Age-related brain changes**

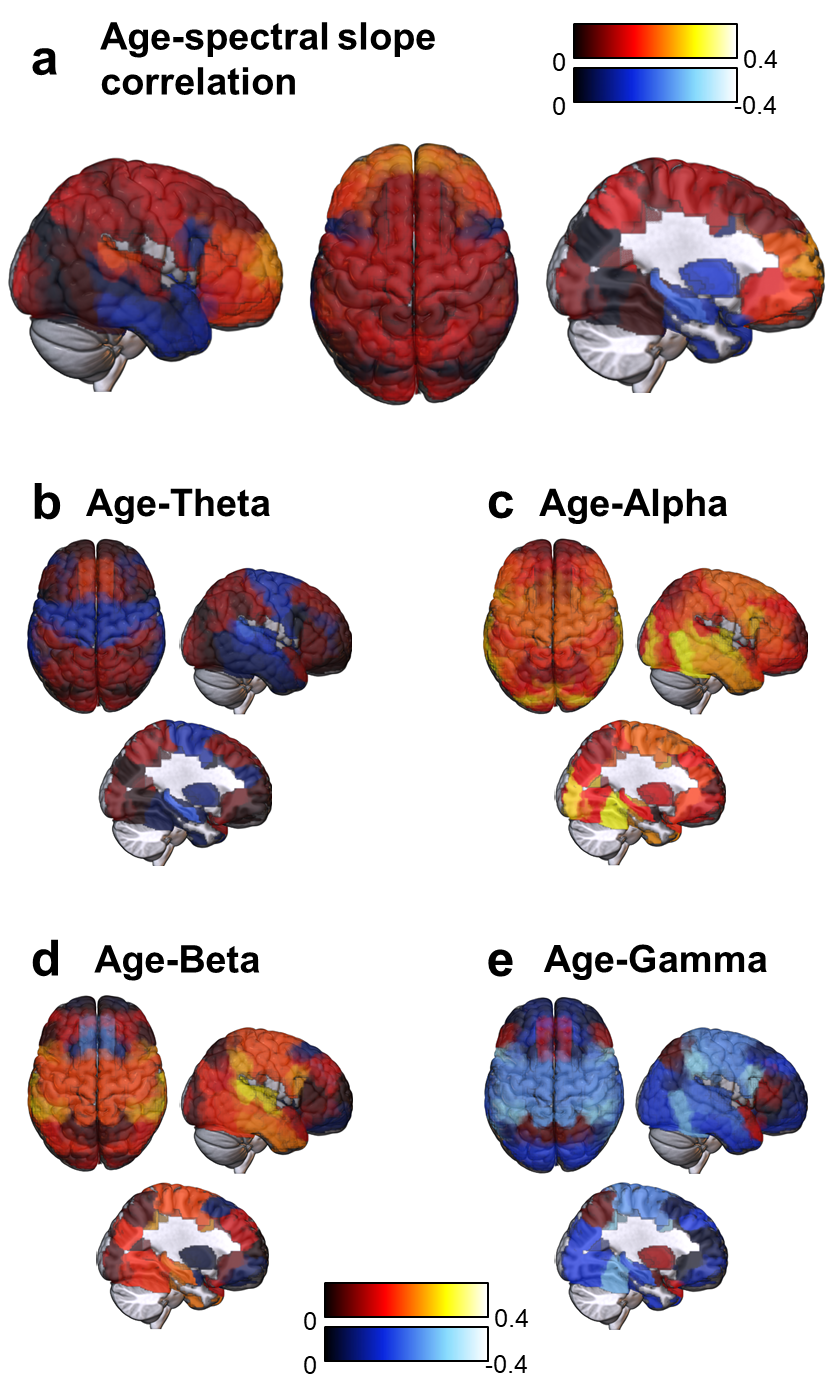

A brain-wide map of correlation between age and spectral slope (**a**, aperiodic), theta (**b**, 3-6Hz), alpha (**c**, 6-10Hz), beta (**d**, 12-18Hz), and gamma (**e**, 40-50Hz) power.

**Supplementary Table 1. Age related brain changes across regions**

| **Region** | **Pearson correlation**  **with age** | | |
| --- | --- | --- | --- |
|  | **r (degree of freedom)** | **p-value (original)** | **p-value**  **(FDR corrected)** |
| **Spectral slope** | | | |
| BA10 | r(39) = 0.434  r(35) = 0.341  r(33) = 0.357  r(39) = -0.354  r(24) = -0.356 | p = 0.005  p = 0.045  p = 0.042  p = 0.023  p = 0.042 | p = 0.133  p = 0.262  p = 0.262  p = 0.262  p = 0.262 |
| BA41&42 |  |  |  |
| BA46 |  |  |  |
| Hippocampus |  |  |  |
| EC&PRC |  |  |  |
| **Theta** | | | |
| BA31 | r(28) = 0.371,  r(35) = -0.348,  r(39) = -0.342, | p = 0.048  p = 0.04  p = 0.029 | p = 0.048  p = 0.04  p = 0.029 |
| BA41&42 |  |  |  |
| Hippocampus |  |  |  |
| **Alpha** | | | |
| BA1&2&3&5 | r(42) = 0.305  r(49) = 0.385  r(30) = 0.401  r(32) = 0.441  r(48) = 0.441  r(57) = 0.47  r(55) = 0.528  r(28) = 0.395  r(42) = 0.574  r(41) = 0.311  r(36) = 0.342  r(51) = 0.356  r(32) = 0.366  r(39) = 0.329 | p = 0.047  p = 0.005  p = 0.023  p = 0.01  p = 0.001  p < 10^-3^  p < 10^-3^  p = 0.034  p < 10^-3^  p = 0.042  p = 0.036  p = 0.01  p = 0.033  p = 0.036 | p = 0.085  p = 0.031  p = 0.066  p = 0.043  p = 0.01  p = 0.002  p < 10^-3^  p = 0.074  p < 10^-3^  p = 0.081  p = 0.074  p = 0.043  p = 0.074  p = 0.074 |
| BA4&6 |  |  |  |
| BA8 |  |  |  |
| BA13 |  |  |  |
| BA20 |  |  |  |
| BA21 |  |  |  |
| BA22 |  |  |  |
| BA31 |  |  |  |
| BA37 |  |  |  |
| BA38 |  |  |  |
| BA39 |  |  |  |
| BA40 |  |  |  |
| BA45 |  |  |  |
| BA47 |  |  |  |
| **Beta** | | | |
| BA1&2&3&5 | r(42) = 0.34  r(49) = 0.341  r(44) = 0.308  r(48) = 0.407  r(57) = 0.361  r(55) = 0.524  r(28) = 0.384  r(24) = -0.496  r(42) = 0.32  r(51) = 0.455  r(35) = 0.537  r(23) = 0.438  r(39) = 0.375 | p = 0.024  p = 0.014  p = 0.037  p = 0.003  p = 0.005  p < 10^-3^  p = 0.044  p = 0.021  p = 0.042  p = 0.001  p = 0.001  p = 0.041  p = 0.016 | p = 0.077  p = 0.065  p = 0.098  p = 0.024  p = 0.029  p < 10^-3^  p = 0.098  p = 0.076  p = 0.098  p = 0.007  p = 0.007  p = 0.098  p = 0.016 |
| BA4&6 |  |  |  |
| BA19 |  |  |  |
| BA20 |  |  |  |
| BA21 |  |  |  |
| BA22 |  |  |  |
| BA31 |  |  |  |
| BA32 |  |  |  |
| BA37 |  |  |  |
| BA40 |  |  |  |
| BA41&42 |  |  |  |
| BA44 |  |  |  |
| Hippocampus |  |  |  |
| **Gamma** | | | |
| BA1&2&3&5 | r(42) = -0.454,  r(49) = -0.424,  r(57) = -0.374,  r(55) = -0.356,  r(42) = -0.508,  r(51) = -0.573,  r(35) = -0.361,  r(23) = -0.601,  r(39) = -0.339, | p = 0.002  p = 0.002  p = 0.005  p = 0.008  p = 0.001  p < 10^-3^  p = 0.033  p = 0.002  p = 0.03 | p = 0.021  p = 0.021  p = 0.021  p = 0.021  p = 0.021  p < 10^-3^  p = 0.053  p = 0.021  p = 0.112 |
| BA4&6 |  |  |  |
| BA21 |  |  |  |
| BA22 |  |  |  |
| BA37 |  |  |  |
| BA40 |  |  |  |
| BA41&42 |  |  |  |
| BA44 |  |  |  |
| Hippocampus |  |  |  |

The FDR correction was based on Benjamini & Hochberg method.

**Supplementary Table 2. The number of participants with electrode implanted in each region.**

| **Region** | **The number of participants** |
| --- | --- |
| BA1&2&3&5 | 44 |
| BA4&6 | 51 |
| BA7 | 31 |
| BA8 | 32 |
| BA9 | 45 |
| BA10 | 41 |
| BA11 | 35 |
| BA13 | 34 |
| BA18 | 22 |
| BA19 | 46 |
| BA20 | 50 |
| BA21 | 59 |
| BA22 | 57 |
| BA31 | 30 |
| BA32 | 26 |
| BA37 | 44 |
| BA38 | 43 |
| BA39 | 38 |
| BA40 | 53 |
| BA41_42 | 37 |
| BA44 | 25 |
| BA45 | 34 |
| BA46 | 35 |
| BA47 | 41 |
| HPC | 41 |
| EC&PRC | 33 |
| Amygdala | 35 |
| Striatum | 33 |

**Supplementary Figure 3.** **Control analyses to verify that aging effects were not associated with epileptic activities**

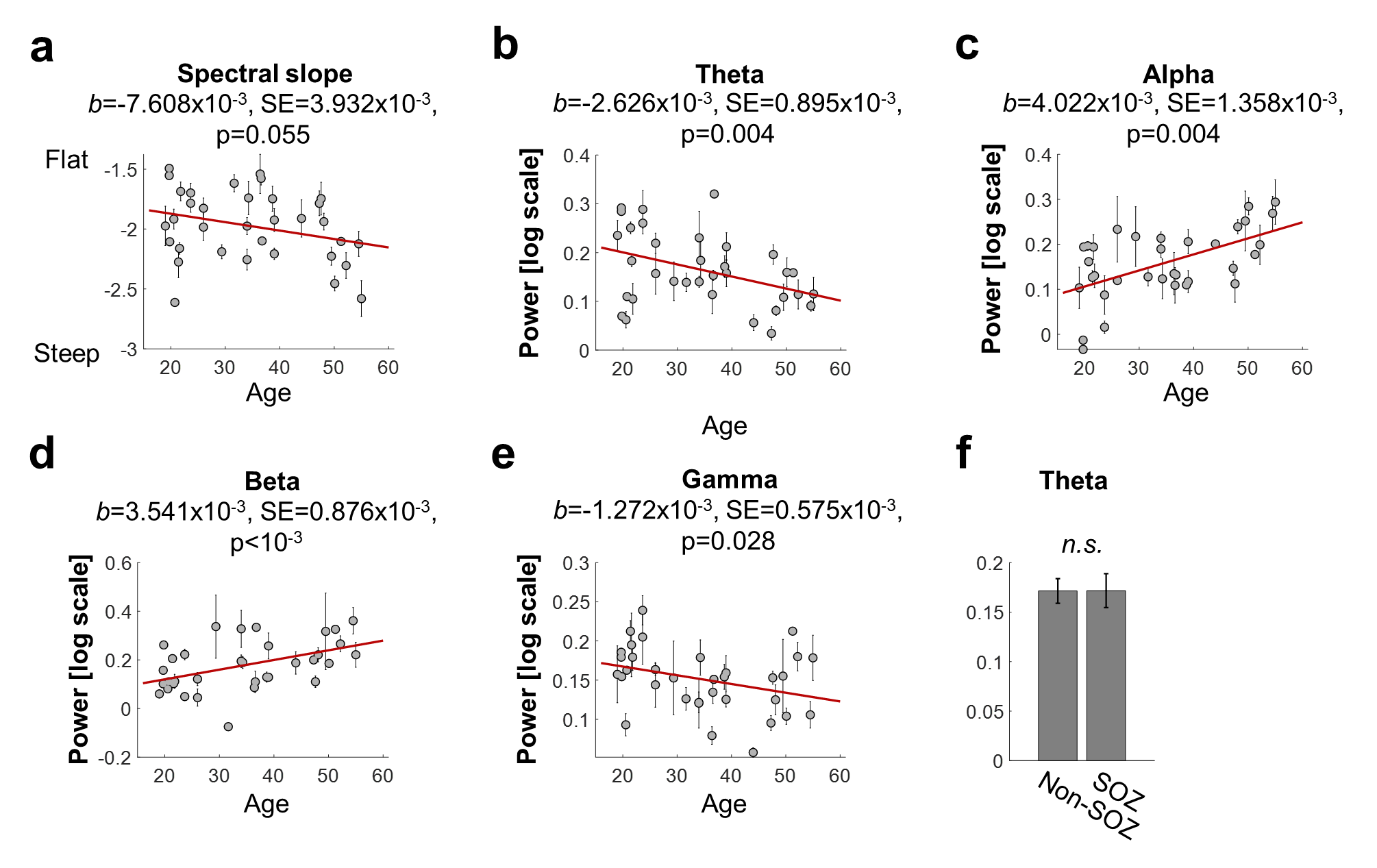

Electrodes in the seizure onset zone (SOZ) were labeled by neurologists based on the daily monitoring of iEEG from all implanted electrodes. To test whether the aging-related hippocampal changes were associated with epileptic neural activities, first we re-applied the same analysis after removing electrodes in the SOZ. **a-e.** Associative patterns between age and isolated periodic band power were still significant. The spectral slope, although it was not statistically significant, showed a trend of being steeper in aging. Therefore, it is unlikely that the epileptic neural activities are a main driver of the age-related hippocampal changes found in this study. **f.** In addition, we compared theta power between electrodes either in (N=21) or outside (N=38) the SOZ. No significant differences were found between electrodes within and outside of the SOZ (t(57)=-0.015 and p=0.988), again supporting the interpretation that age-related changes can be dissociated from epileptic neural activity.

**Supplementary Figures 4-5 and Tables 3-5. Control analyses to verify that aging effects are dissociable from epilepsy duration and education length**

To assess whether age-related changes in hippocampal markers were confounded by education length or epilepsy duration, we performed control analyses using an independent dataset (Dataset 2). This dataset comprises intracranial EEG recordings from 46 bipolar electrode pairs implanted in the hippocampus of 27 epilepsy patients at Seoul National University Hospital. Education level was reported by 15 patients (9, 12, or 16 years—the standard durations of middle, high schools or university based on Korean education system), epilepsy duration was available for 24 patients, and information of anti-epileptic drug intake was available for 26 patients. All participants were on daily anti-epileptic medication during the iEEG recordings. Due to the limited number of cases, patients with comorbid neurological disorders were excluded from the present analyses.

Recordings were obtained during awake resting state, with durations ranging from 1 to 5 minutes across individuals. We applied spectral parameterization (formally referred as “FOOOF”) to the 2–50 Hz frequency range, and extracted the spectral slope and band power (after flattening) in the theta (3–6 Hz), alpha (6–10 Hz), beta (12–18 Hz), and gamma (40–50 Hz) bands, consistent with the main analyses.

**(1) Reproduction of age-related patterns in hippocampal aperiodic and periodic components**

**Supplementary Figure 4. a.** A histogram for number of patients by age. **b-f.** Scatter plots for hippocampal aperiodic and periodic components by age.

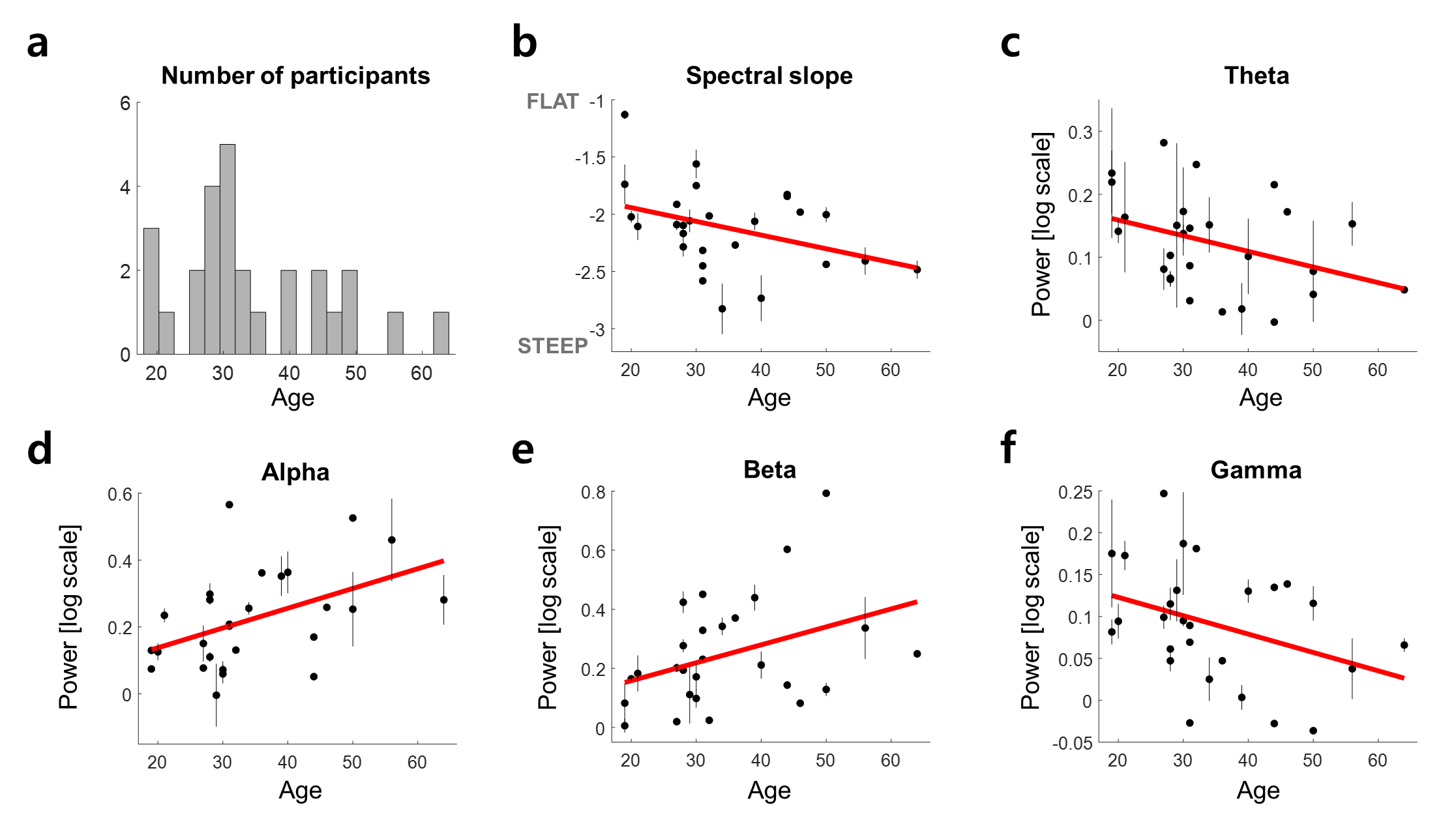

**Supplementary Table 3.** Comparison of linear mixed-effects model results assessing the relationship between hippocampal aperiodic/periodic components and age in the original and new datasets.

| Linear mixed-effects models | **Original dataset 1**  **(spatial memory navigation and encoding)** | **Dataset 2**  **(resting state)** |
| --- | --- | --- |
| Spectral slope | b=-7.672x10^-3^, SE=3.56x10^-3^, **p=0.032** | b=-12.481x10^-3^, SE=5.75x10^-3^, **p=0.035** |
| Theta | b=-2.123x10^-3^, SE=0.866x10^-3^, **p=0.015** | b=-2.02x10^-3^, SE=0.979x10^-3^, **p=0.045** |
| Alpha | b=2.632x10^-3^, SE=1.182x10^-3^, **p=0.027** | b=6.004x10^-3^, SE=2.035x10^-3^, **p=0.005** |
| Beta | b=1.902x10^-3^, SE=0.949x10^-3^, **p=0.046** | b=5.91x10^-3^, SE=2.756x10^-3^, **p=0.038** |
| Gamma | b=-1.029x10^-3^, SE=0.493x10^-3^, **p=0.038** | b=-2.108x10^-3^, SE=1.026x10^-3^, **p=0.046** |

First, we examined age-related patterns in aperiodic (i.e., spectral slope) and periodic components (i.e., theta, alpha, beta, and gamma power) using Dataset 2 and compared the results with those obtained from Dataset 1. Consistent with the main analyses, we implemented linear mixed-effects models, with age as a predictor, resting-state aperiodic/periodic components as dependent variables, and individual participants included as a random-effect variable for grouping electrodes (i.e., aperiodic/periodic components ~ age + 1|participant).

The statistical results (see figure and table above) indicated that age-related effects in Dataset 2 replicated those in Dataset 1: (1) steeper spectral slope (**b**), (2) decreased theta and gamma power (**c and f**), and (3) increased alpha and beta power with age (**d and e**).

**(2) Effects of epilepsy duration and education level on hippocampal aperiodic and periodic components**

Next, we investigated whether hippocampal spectral features were influenced by epilepsy duration and education level. We first assessed the effect of epilepsy duration using linear mixed-effects models, with epilepsy duration as a predictor, aperiodic/periodic components during resting state as dependent variables, and individual participants as a random-effect variable for grouping electrodes (i.e. aperiodic/periodic components ~ duration of epilepsy + 1|participant). To assess the effect of education, a separate linear mixed-effects model was used: aperiodic/periodic components ~ education length + 1|participant. Similarly, the effect of the number of anti-epilepsy drug intake was identified by another linear mixed-effects model: aperiodic/periodic components ~ the number of anti-epilepsy drug intake + 1|participant.

**Supplementary Figure 5.** Scatter plots for hippocampal aperiodic and periodic components by duration of epilepsy (**a-e**), education length (**f-j**), or the number of anti-epileptic drug intake (**k-o**).

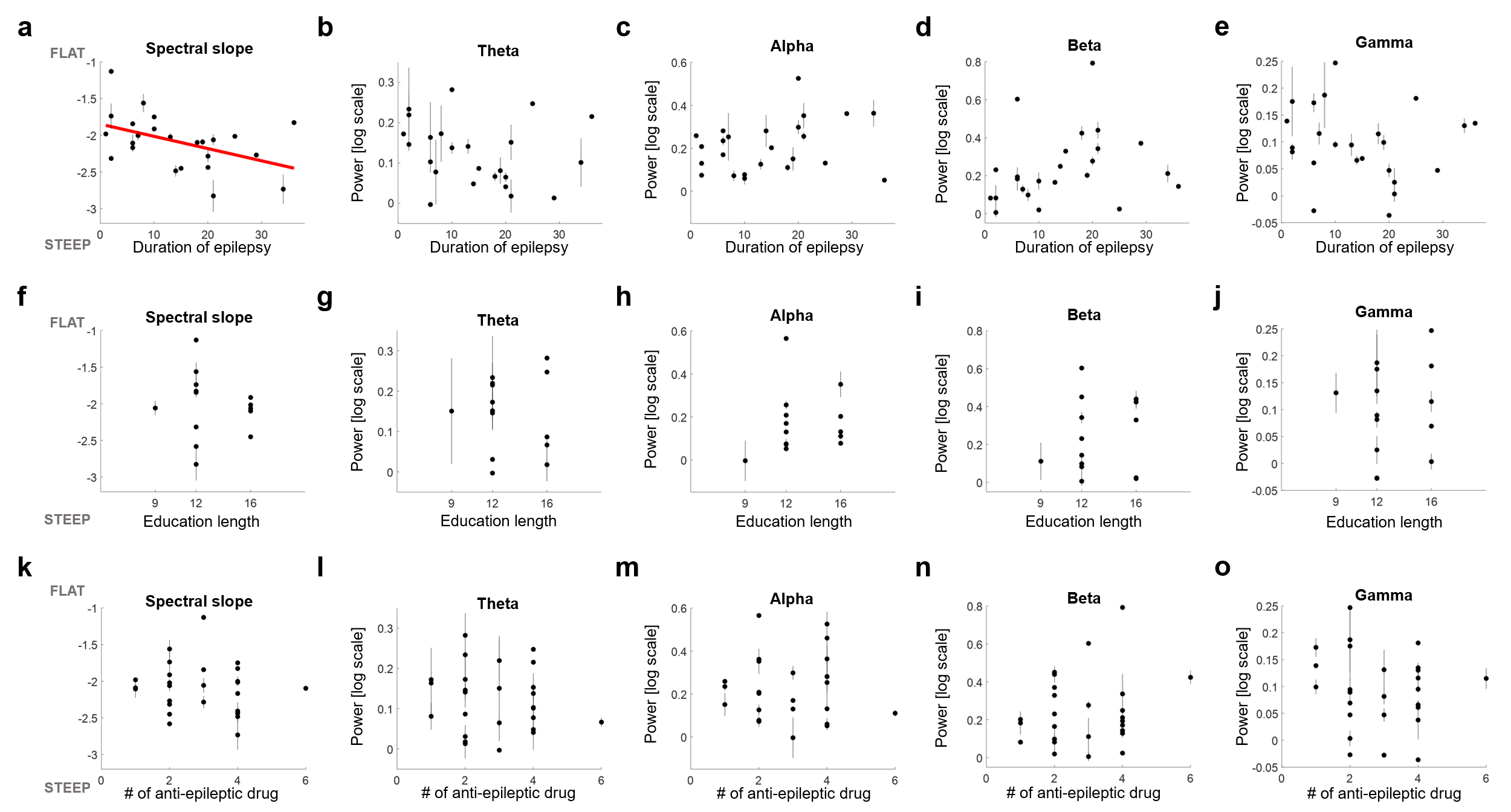

**Supplementary Table 4.** Results of linear mixed-effects models assessing the relationship between hippocampal aperiodic/periodic components and patient-specific factors.

| Linear mixed-effects models | **Duration of epilepsy** | **Education length** | **Anti-epileptic drug** |
| --- | --- | --- | --- |
| Spectral slope | b=-18.438x10^-3^, SE=6.814x10^-3^, **p=0.01** | b=-21.761x10^-3^, SE=51.557x10^-3^, p=0.677 | b=-31.774x10^-3^, SE=54.2x10^-3^, p=0.561 |
| Theta | b=-2.613x10^-3^, SE=1.456x10^-3^, p=0.08 | b=-11.415x10^-3^, SE=9.546x10^-3^, p=0.244 | b=-8.384x10^-3^, SE=11.391x10^-3^, p=0.466 |
| Alpha | b=3.597x10^-3^, SE=2.264x10^-3^, p=0.12 | b=10.622x10^-3^, SE=16.328x10^-3^, p=0.522 | b=3.232x10^-3^, SE=22.998x10^-3^, p=0.889 |
| Beta | b=4.731x10^-3^, SE=3.604x10^-3^, p=0.197 | b=11.076x10^-3^, SE=22.085x10^-3^, p=0.621 | b=29.23x10^-3^, SE=27.906x10^-3^, p=0.301 |
| Gamma | b=-1.302x10^-3^, SE=1.343x10^-3^, p=0.338 | b=2.123x10^-3^, SE=9.531x10^-3^, p=0.826 | b=-7.621x10^-3^, SE=10.331x10^-3^, p=0.465 |

The linear mixed-effects model revealed that most hippocampal neural components were not significantly associated with epilepsy duration (**b–e**), with the exception of the spectral slope, which showed a significant negative relationship (**a**). There was a strong negative trend between hippocampal theta power and epilepsy duration (**b**), but it was not statistically significant. In contrast, none of the hippocampal aperiodic or periodic components were significantly associated with education length, as assessed by linear mixed-effects modeling (**f–j**).

**(3) Dissociating the effects of epilepsy duration and education length from age-related changes**

To determine whether the observed relationship between spectral slope and epilepsy duration was independent of age, we performed partial correlation analyses controlling for age. After accounting for age, the association between spectral slope and epilepsy duration became not significant but remained at a strong trend level (r=-0.404, p=0.056; *r*=-0.457, *p*=0.025 without accounting for age). This suggests that a steeper hippocampal spectral slope may be influenced by both aging and longer epilepsy duration. In contrast, the negative correlation between theta power and epilepsy duration became weaker after controlling for age, indicating that changes in theta power were more closely related to aging than to epilepsy history.

No hippocampal neural components were significantly associated with education level after controlling for age via partial correlation. However, this analysis may have been underpowered due to the limited number of patients who reported their education level.

**Supplementary Table 5.** Results of partial correlation controlling for age to assess the relationship between hippocampal aperiodic/periodic components and patient-specific factors.

| Partial correlation | **Duration of epilepsy** | **Education length** | **Anti-epileptic drug** |
| --- | --- | --- | --- |
| Spectral slope | r=-0.404, p=0.056 | r=-0.107, p=0.727 | r=0.050, p=0.811 |
| Theta | r=-0.106, p=0.630 | r=-0.020, p=0.949 | r=0.026, p=0.903 |
| Alpha | r=0.166, p=0.449 | r=0.131, p=0.670 | r=-0.179, p=0.392 |
| Beta | r=0.169, p=0.442 | r=0.097, p=0.751 | r=0.069, p=0.745 |
| Gamma | r=-0.057, p=0.795 | r=0.191, p=0.532 | r=-0.005, p=0.980 |

In summary, the marginal influence of epilepsy duration and education level—aside from the spectral slope—substantiates our main conclusion that changes in hippocampal aperiodic and periodic components are primarily age-dependent. Notably, these control analyses further support the robustness of our key finding that reduced hippocampal theta and gamma power underlies age-related spatial memory decline.

**Supplementary Figure 6. Lateralization in the hippocampus and DLPFC**

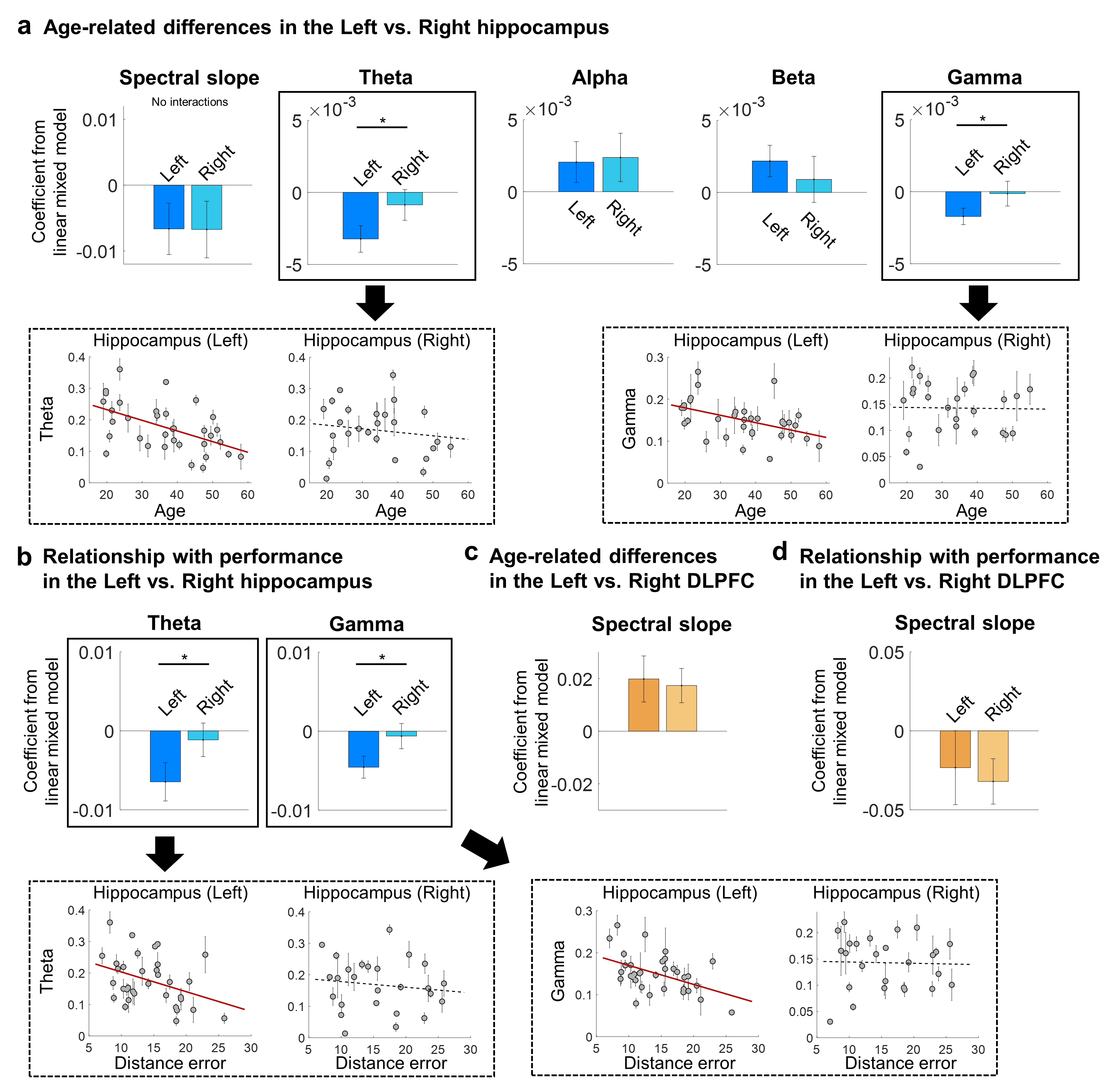

With the main dataset, we implemented linear mixed models to compare aging-related changes between the left and right hippocampus. **a.** Lateralization of aging effects was found in hippocampal theta and gamma power (theta: F(1,217)=3.89, p=0.049; gamma: F(1,217)=4.723, p=0.031) but not in spectral slope, alpha, and beta power (spectral slope: F(1,217)=0.111, p=0.74; alpha: F(1,217)=0.486, p=0.487; beta: F(1,217)=1.1, p=0.295). The age-related decline of theta and gamma power was found only in the left hippocampus (theta: b=-3.232x10^-3^, SE=0.92x10^-3^, p<10^-3^; gamma: b=-1.718x10^-3^, SE=0.567x10^-3^, p=0.003; Right hippocampus theta: b=-0.867x10^-3^, SE=1.068x10^-3^, p=0.418; gamma: b=-0.133x10^-3^, SE=0.861x10^-3^, p=0.878). **b.** Similar to the aging effect, lateralization was found in a relationship between theta power and spatial memory performance (theta: F(1,217)=7.479, p=0.0068; gamma: F(1,217)=5.269, p=0.023). Again, a negative relationship between theta/gamma power and distance error was only found in the left hippocampus (theta: b=-6.459x10^-3^, SE=2.429x10^-3^, p=0.009; gamma: b=-4.564x10^-3^, SE=1.395x10^-3^, p=0.001; Right hippocampus theta: b=-1.144x10^-3^, SE=2.124x10^-3^, p=0.591; gamma: b=-0.623x10^-3^, SE=1.601x10^-3^, p=0.698). **c-d.** Lastly, we found no significant differences across left and right hemispheres in a relationship between DLPFC spectral slope and age (**c**) or performance (**d**) (aging: F(1,113)=0.001, p=0.991; performance: F(1,113)=0.087, p=0.769).

**Supplementary Figure 7. Original power spectrum**

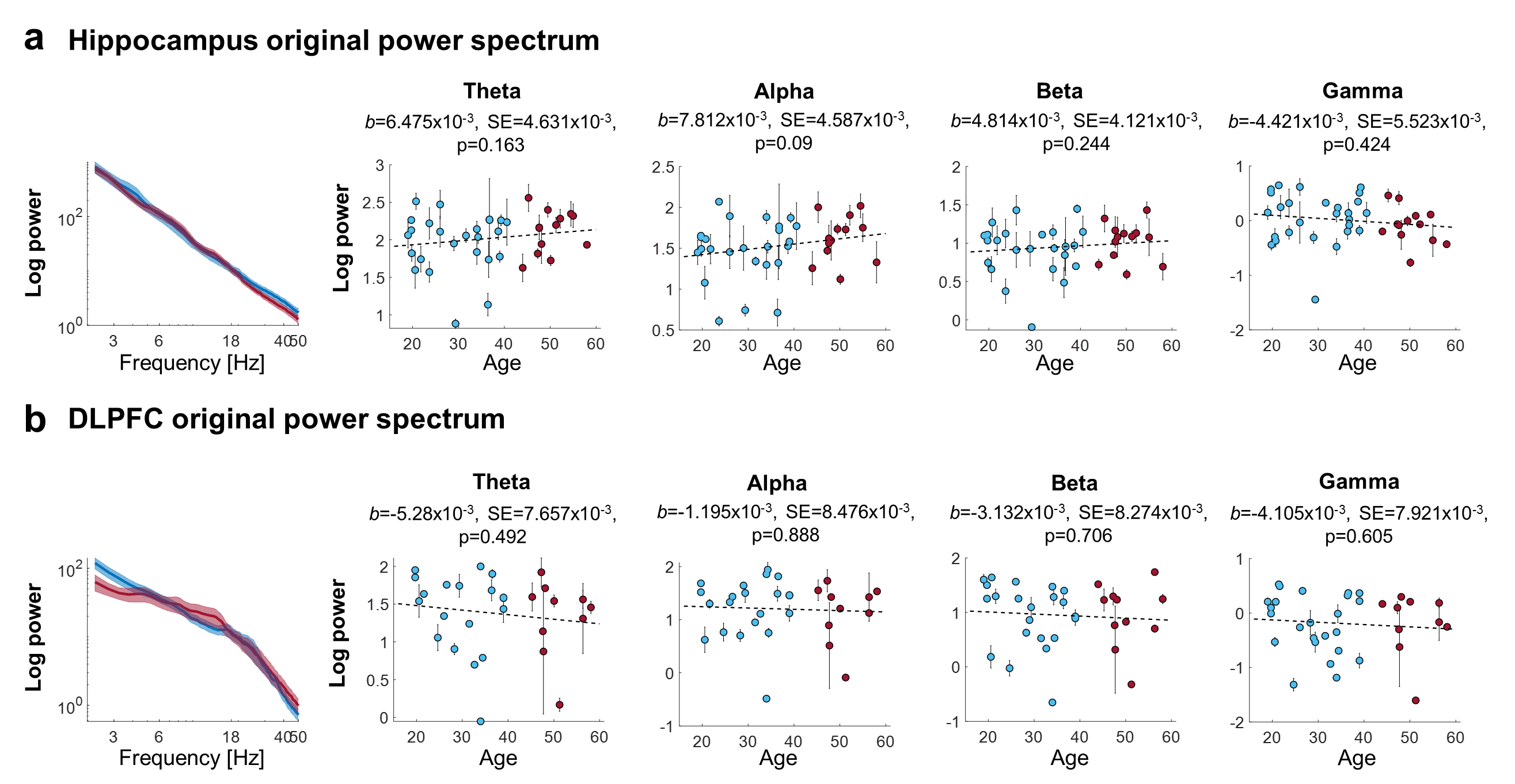

**a.** Unlike the strong association between isolated band power and age (Fig. 2i), band power from the raw power spectrum (i.e. before subtracting the spectral fitting line) did not show a clear increase or decrease in aging. **b.** Similarly, no relationship between age and band power from the raw spectrum was observed in the DLPFC. These results emphasize the importance of dissociating aperiodic and periodic components for a more precise characterization of iEEG spectral features.

**Supplementary Figure 8. Behavioral metrics of spatial memory performance**

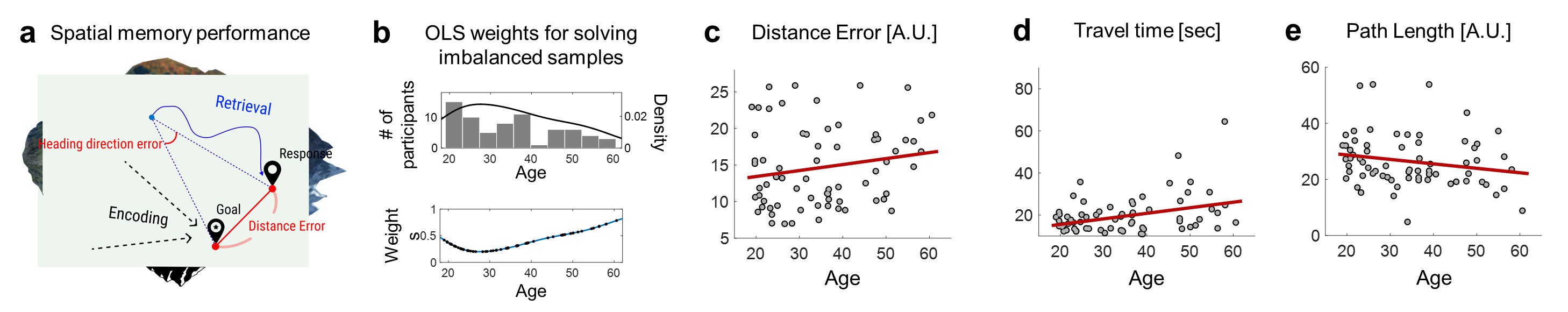

**a.** A schematic of the spatial memory task and behavioral metrics. **b.** We compared behavioral metrics across the whole age range by implementing general linear models with the observation weights proportional to the age to balance out the lower sample size of the older participants. (top) Kernel density estimation with the Gaussian kernel was used to calculate the probabilistic density of age, which was shown together with the age histogram. (bottom) Weights (blue line) were calculated by $1-\alpha\times density, \alpha=28.916$ where the hyperparameter $\alpha$ was decided to make weights ranging from 0.2 to 0.8. The black dots indicate the actual weights used for our samples. **c.** Distance error represented how far the participants’ response was from the actual target location, and we found a higher error in the older participants (b=0.111, SE=0.051, p=0.034). d**.** Travel time was longer for the old participants (b=0.296, SE=0.097, p=0.003). **e.** Path length during retrieval showed a trend of decrease in aging (b=-0.16, SE=0.093, p=0.09). Overall, older participants showed higher distance errors and required a longer exploration period during the retrieval.

**Supplementary Figure 9. Exploring performance markers in the hippocampus and DLPFC**

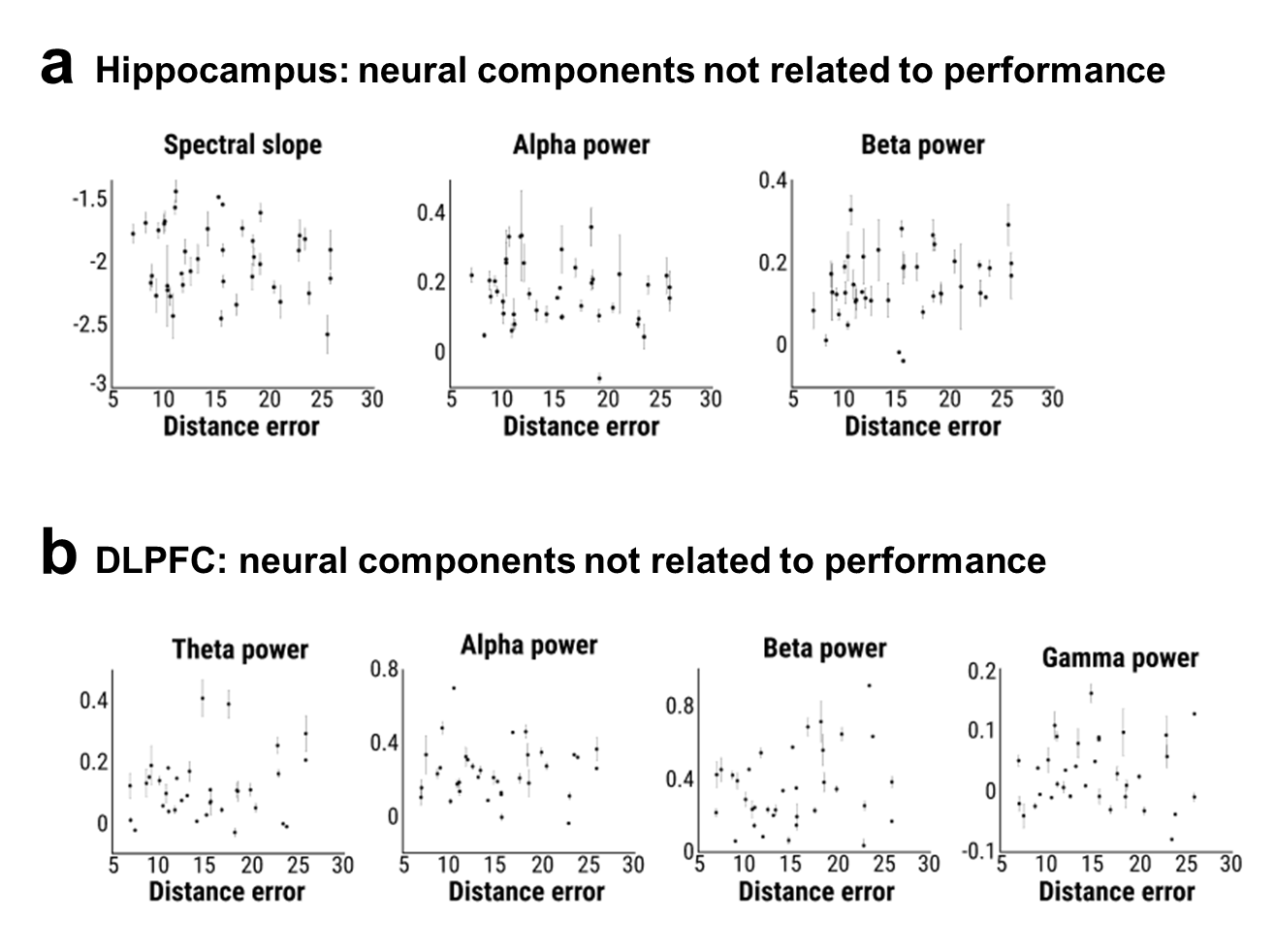

**a.** Neural components in the hippocampus that are not related to distance error based on linear mixed models (spectral slope: b=-8.695x10^‑3^, SE=7.856 x10^‑3^, p=0.27; alpha: b=-2.304x10^‑3^, SE=2.656x10^‑3^, p=0.387; beta: b=3.92x10^‑3^, SE=2.026x10^‑3^, p=0.054). **b.** Neural components in the DLPFC which are not related with distance error based on linear mixed models (theta: b=3.698x10^‑3^, SE=3.179x10^‑3^, p=0.247; alpha: b=0.567x10^‑3^, SE=4.349x10^‑3^, p=0.896; beta: b=8.78x10^‑3^, SE=6.121x10^‑3^, p=0.154; gamma: b=0.304x10^‑3^, SE=1.633x10^‑3^, p=0.853).

**Supplementary Figure 10. Hippocampal subregions and medial temporal lobe (MTL)**

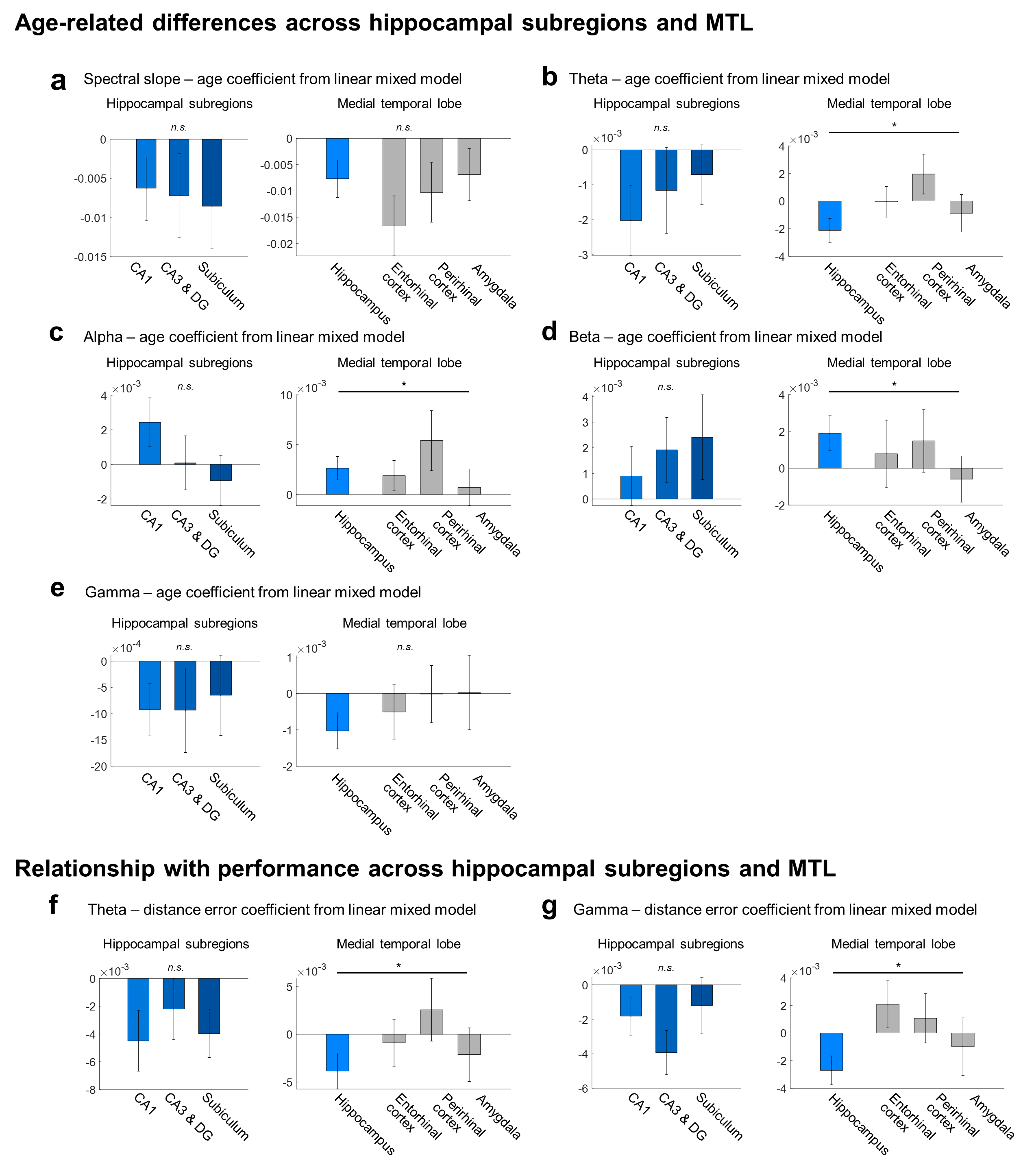

We measured and compared age-related differences or a relationship with performance across the hippocampal subregions and medial temporal lobe by implementing linear mixed models. **a.** Aging effects in spectral slope were not significantly different across the hippocampal subregions (age-region interaction: F(2,208)=1.687, p=0.188) and the medial temporal lobe (F(3,444)=1.942, p=0.122). **b.** Aging effects in theta power were not significantly different across the hippocampal subregions (F(2,207)=0.418, p=0.659). However, age-related theta decrease was significantly pronounced in the hippocampus compared to the other medial temporal lobe regions (age-region interaction: F(3,440)=7.99, p<10^-3^). **c-d.** Similar to theta, age-related increases in alpha and beta power were significantly pronounced in the hippocampus compared to the other medial temporal regions (alpha: F(3,444)=3.753, p=0.011; beta: F(3,443)=4.065, p=0.007) despite that the perirhinal cortex also showed a pronounced age-related alpha power increase. Again, age-related differences were similar across the hippocampal subregions (alpha: F(2,203)=2.278, p=0.105; beta: F(2,208)=1.390, p=0.251). **e.** Age-related decreases in gamma power were not significantly different across the hippocampal subregions (F(2,208)=0.209, p=0.812) and the medial temporal lobe (F(3,444)=1.45, p=0.228). **f-g.** A relationship between distance error and theta and gamma power was significantly weaker in the other medial temporal lobe regions compared to the hippocampus (distance error-region interaction theta: F(3,440)=3.657, p=0.013; gamma: F(3,444)=3.681, p=0.012). However, a relationship with performance was not different across hippocampal subregions (theta: F(2,207)=0.45, p=0.638; gamma: F(2,208)=0.856, p=0.426).

**Supplementary Figure 11. Further analyses of retrieval in the Match and Nonmatch conditions**

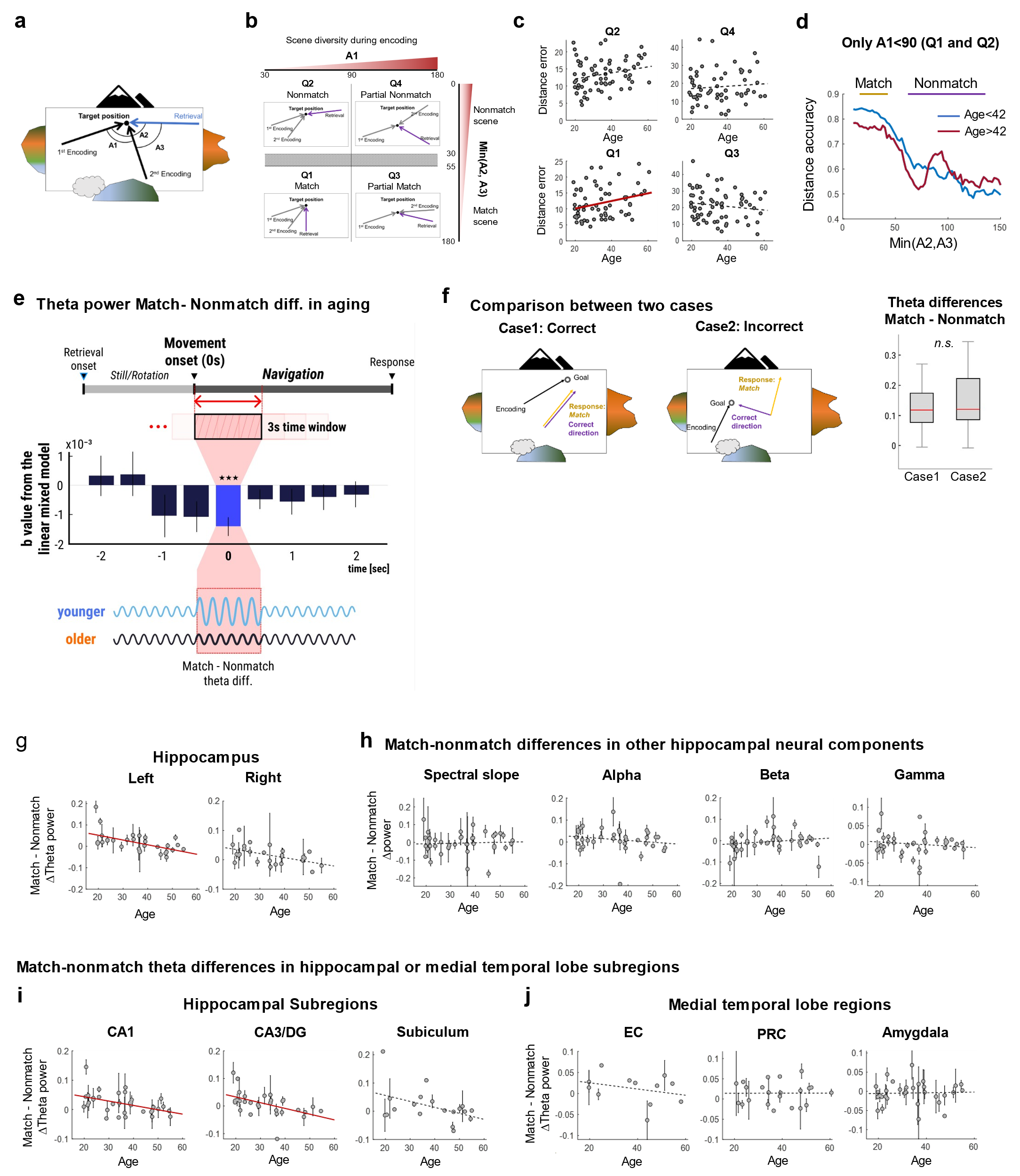

**a.** A schematic of the environment (bird’s-eye view) with an example of the two encoding paths and the retrieval path. A1, A2, and A3 represent angles between two encoding paths (A1) or between the retrieval and each of the encoding paths (A2 and A3). **b.** Trials were divided into four conditions based on A1, A2, and A3. Q1(Match): A1<90 & min(A2, A3)<30. Q2(Nonmatch): A1<90 & min(A2, A3)>55. Q3(Partial Match): A1>90 & min(A2, A3)<30. Q4(Partial Nonmatch): A1>90 & min(A2, A3)>55. (min(A2, A3) indicates the smaller value between A2 and A3). **c.** Age-related spatial memory decline was only shown in the Q1 condition (b=0.12, SE=0.049, p=0.016) while distance error did not show any relationship with age (Q2: b=-0.107, SE=0.09, p=0.239; Q3: b=0.088, SE=0.044 p=0.051; Q4: b=0.066, SE=0.098, p=0.5). **d.** We measured the distance accuracy across min(A2, A3) when A1<90 and found clearly different patterns between min(A2, A3) below 30 and above 55 degrees, as well as a transition zone between the two angles. **e.** Top: The relationship between age and theta power difference (i.e. *Match* - *Nonmatch*), measured by linear mixed-effects models, using a 3s moving window (moving step=0.5s). Bottom: A schematic diagram of hippocampal theta power during the retrieval phase, from retrieval onset, to movement onset, to the response. The age-related difference in the landmark-dependent theta engagement was more pronounced in the first three seconds of navigation onset, which is consistent with the interpretation that participants used the scene for deciding their orientation to the target object (FDR-corrected p<10^-3^). **f.** In Cases 1 and 2, we compared participants who navigated in a particular direction with respect to the Match view, across trials in which the correct answer was the Match (Case 1) or Nonmatch (Case 2) view by implementing linear mixed models with age and condition as a predictor. Theta power decreased in aging but did not show any difference between Case 1 and Case 2 (condition effect: F(1,368)=0.092, p=0.762), suggesting that the increased theta in the Match view was not simply attributable to better performance. **g.** The age-related difference in theta engagement in the Match condition was pronounced in the left hippocampus (b=-2.064x10^-3^, SE=0.553x10^-3^, p=0.003), but the right hippocampus also showed a trend (b=-1.031x10^-3^, SE=0.524x10^-3^, p=0.052). An interaction between hemispheres (i.e. left and right) and age was not significant. **h.** Hippocampal spectral slope, alpha, beta, and gamma power were not associated with the age-related Match/Nonmatch conditional differences (spectral slope: b=0.619x10^-3^, SE=0.692x10^-3^, p=0.372; alpha: b=-0.695x10^-3^, SE=0.563x10^-3^, p=0.219; beta: b=0.509x10^-3^, SE=0.625x10^-3^, p=0.416; gamma: b=-0.264x10^-3^, SE=0.374x10^-3^, p=0.481). **i.** The age-related difference in theta engagement in the Match condition was pronounced in CA1 and CA3/DG (CA1: b=-1.1602x10^-3^, SE=0.449x10^-3^, p=0.011; CA3/DG: b=-2.316x10^-3^, SE=0.649 x10^-3^, p<10^-3^), but the subiculum also showed a trend (b=-1.78x10^-3^, SE=0.873x10^-3^, p=0.052). **j.** Unlike in the hippocampus, Match-Nonmatch theta power differences were not significantly associated with age in the other medial temporal lobe regions (age-region interaction: F(3,403)=2.942, p=0.033; entorhinal cortex: b=-0.609x10^-3^, SE=0.783x10^-3^, p=0.445; perirhinal cortex: b=0.138x10^-3^, SE=0.396x10^-3^, p=0.729; amygdala: b=0.033x10^-3^, SE=0.567x10^-3^, p=0.954).
